# Anisotropic Cell Shape and Motion Coordinate Hindbrain Neuropore Morphogenesis

**DOI:** 10.1101/2024.11.21.624679

**Authors:** Fernanda Pérez-Verdugo, Eirini Maniou, Gabriel L. Galea, Shiladitya Banerjee

## Abstract

Neural tube closure is a critical morphogenetic process in vertebrate development, and failure to close cranial regions such as the hindbrain neuropore (HNP) leads to severe congenital malformations. While mechanical forces like actomyosin purse-string contraction and directional cell crawling have been implicated in driving HNP closure, how these forces organize local cell shape and motion to produce large-scale tissue remodeling remains poorly understood. Using live and fixed imaging of mouse embryos combined with cell-based biophysical modeling, we show that these force-generating mechanisms are insufficient to explain the robust patterns of cell elongation and nematic alignment observed at the HNP border. Instead, we show that local anisotropic stress and cytoskeletal organization are required to generate these patterns and promote midline cell motion. Our model captures key features of cell shape dynamics and emergent nematic order, which we confirm experimentally, including the alignment of actin fibers with cell shape and enhanced midline cell speed. Comparative analysis with chick embryos, which lack supracellular purse-strings, supports a conserved link between tension generation and cellular patterning. These findings establish a physical framework connecting force generation, cell shape anisotropy, and tissue morphodynamics during epithelial gap closure.

---

Epithelial gap closure is a complex process crucial for morphogenesis during early development and essential for maintaining the integrity and functionality of epithelial barriers throughout the lifespan of multicellular organisms [1, 2]. Gap closure spans a wide range of length scales, ranging from the closure of single-cell-sized gaps during apoptosis [3] to the closure of gaps of the size of hundreds of cells during morphogenesis [4–6]. Previous work has identified two key force-generating mechanisms driving gap closure in many systems: cell crawling and actomyosin purse-string contraction [6–10]. However, how these distinct mechanisms are physically coordinated across space and time for gap closure *in vivo* remains unknown. In particular, it is poorly understood how local cellular forces give rise to the emergent patterns of cellular organization to coordinate tissue shape changes.

During vertebrate development, the neural tube forms the central nervous system through a series of tissue folding and fusion events. Failure to close the neural tube leads to severe congenital disorders such as spina bifida [11] and anencephaly [12], affecting nearly 300,000 births annually [13]. In mammals, the final step in cranial neural tube closure is closure of the hindbrain neuropore (HNP) – a large epithelial gap that seals through progressive “zippering” from both ends [14, 15]. Mouse embryos exhibit multi-site closure dynamics similar to humans, making them a relevant model for studying the mechanics of neuropore closure [16, 17].

We previously showed that HNP closure depends critically on the surface ectoderm (future epidermis), which bridges the gap by extending into the neuropore and forming stable contact points across the embryonic midline. We identified two major force-generating mechanisms in the surface ectoderm driving HNP closure: supracellular actomyosin purse-string contraction along the gap edge, and directional cell crawling toward the midline [6]. While these mechanisms explain bulk tissue movement during HNP closure, they do not account for a striking and reproducible feature: cells at the leading edge of the gap exhibit anisotropic shapes aligned along the rostrocaudal axis, forming a narrow band of nematically ordered morphology (Fig. **1**A). These cells also move more rapidly during closure, suggesting a link between cell shape, directional motion, and tissue-scale morphogenesis. This raises key questions: How does spatial patterning of cell shape and nematic order emerge in this system? Can these patterns arise through intrinsic forces rather than pre-patterned cues? And what role do they play in coordinating force generation and tissue remodeling during gap closure? Prior theoretical work has identified nematic alignment and mechanical feedback as drivers of collective behavior in tissues [18, 19], but these ideas have not been directly tested in a well-defined *in vivo* context.

**Fig. 1.**
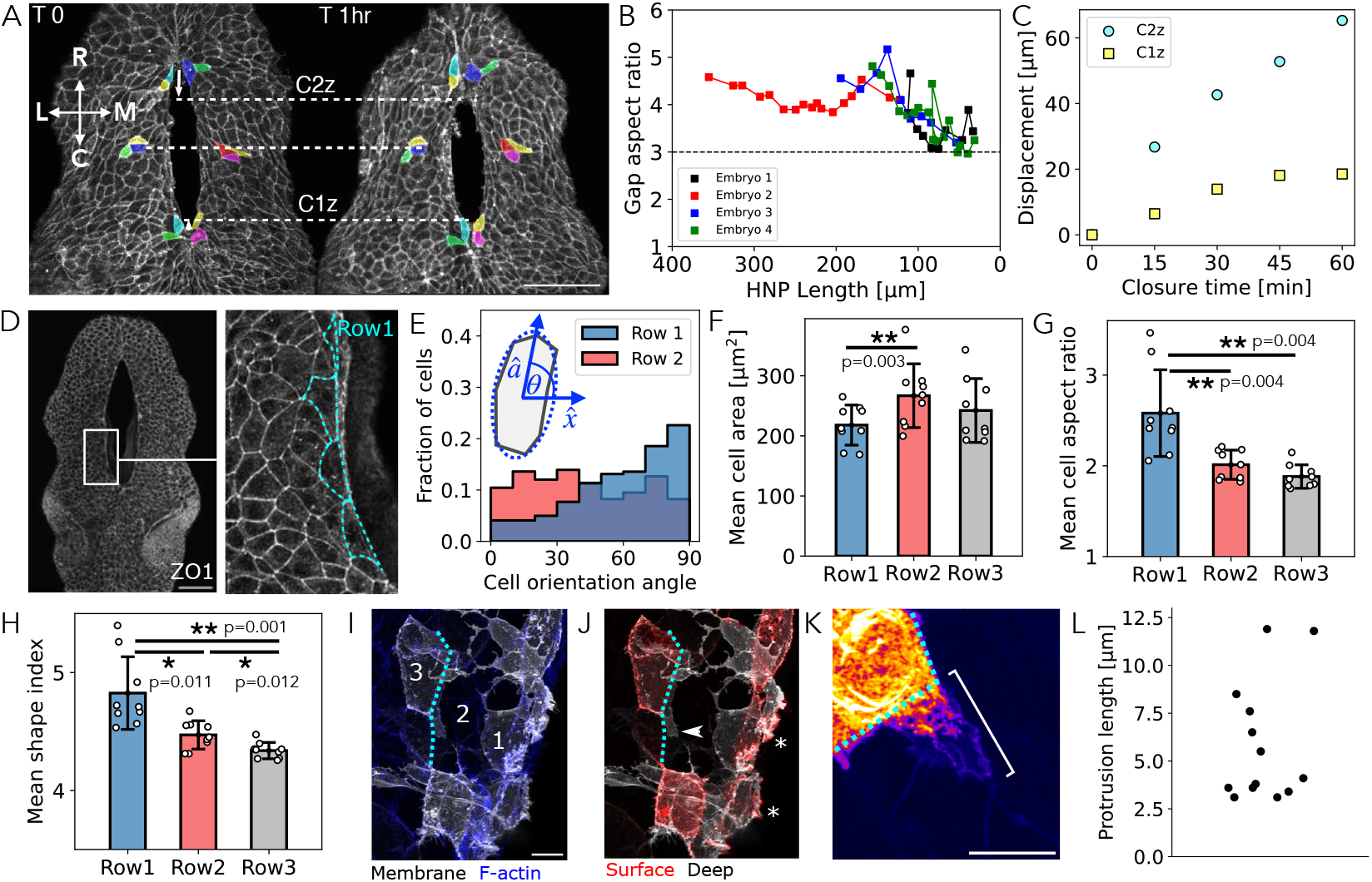
Surface ectoderm cells exhibit robust shape patterning during hindbrain neuropore closure. (A) HNP gap in a 14 somite live-imaged mouse embryo, illustrating the asymmetric closure and the shape evolution of individual cells. C1z and C2z indicate the zippering points progressing from Closure points 1 and 2. Dashed lines indicate landmark positions after 1 hour of live imaging. Colors indicate individual cells. Note cells in contact with C2z at T0 have lost contact with it one hour later whereas cells at C1z remain in contact for the full hour. Scale bar, 100 µm. The solid white arrows indicate the rostro-caudal (R-C) and medio-lateral (M-L) axes. (B) Gap aspect ratio (height over width) versus HNP length, for n=4 live-imaged embryos. Embryos 1-3 (15-16 somite) were previously published [6], and Embryo 4 is shown in (A). (C) Quantification of C2z and C1z (zippering points) displacement over time (from *T* = 0, Fig. 1A). (D) HNP gap in a 15 somite fixed embryo, highlighting the elongation of row-1 surface ectoderm cells (cyan) along the gap. Scale bar, 100 µm. (E-H) Shape analysis of cells in Row 1 (n = 221), Row 2 (n = 316) and Row 3 (n = 376) from 9 embryos. (E) Row-1 and row-2 cellular orientation distributions. The normal vector **â** defines the cell orientation direction. Cell orientation angle is defined such that *θ* = 90^°^ indicates elongation along the rostro-caudal axis. (F) Mean cell area per row. (G) Mean cell aspect ratio per row. (H) Mean cell shape index per row. The statistical analysis shown in (F-G) considers individual embryos (n=9) as the unit of measure. Individual data points are represented with white dots. * *P <* 0.05, * * *P <* 0.01. The analysis shown in (E-H) excludes cells whose centers lie above C2z and below C1z. Error bars represent ±1 standard deviation. (I) Mosaic cell membrane label (BioT) counter-stained with phalloidin to visualize F-actin. (J) Surface subtraction of BioT-labelled cells at the HNP margin showing the apical surface (red, corresponding to the polygons modelled in simulations) and deeper structures. Row 1 cells extend ruffles into the gap (*). Arrows indicate a cryptic lamellipodium-like protrusion extending beyond the apical cell cortex of a Row 3 cell (dashed line) underneath a Row 2 cell ahead of it. (K) Higher magnification of a cryptic lamellipodium-like protrusion indicating its length (white bracket) ahead of the cell cortex (dashed line). Scale bars, 10 µm. (L) Quantification of protrusion lengths of Row 2-3 cells (13 protrusions from 4 embryos with 14-15 somites).

Here, we address these question using a combination of high-resolution imaging and biophysical modeling of HNP closure in mouse embryos. We mapped cell shapes and orientations during HNP closure and uncovered significant heterogeneities in cell shape and orientation across the surface ectoderm. These patterns—particularly the rostrocaudal alignment and elongation of cells near the gap—are highly reproducible across embryos. Such spatial organization could either be pre-specified by genetic or signaling cues, or may arise from mechanical feedback within the tissue.

To test this, we developed a biophysical model of HNP closure incorporating both cell crawling and supracellular purse-string contraction. While sufficient to reproduce tissue-scale closure dynamics, this model fails to account for the observed cell shape and orientation patterning. We show that adding a mechanosensitive feedback loop – where local shear stress aligns cytoskeletal elements and reinforces cell elongation – generates robust, spatially patterned cell morphologies consistent with experimental data. In this model, purse-string tension acts as a mechanical cue that triggers local cytoskeletal alignment, leading to rostrocaudal elongation near the gap and the emergence of a midline region with enhanced cell motion. In support of this mechanism, we find that chick embryos, which lack actomyosin purse-strings during neuropore closure, also lack rostrocaudally elongated surface ectoderm cells. Together, our results reveal that mechanical feedback between cell shape and internal stress organization can generate patterned anisotropy to guide epithelial gap closure.

## Robust cell shape patterning during hindbrain neuropore closure

The HNP gap closure process exhibits robust tissue-level geometric features [6], including the maintenance of an elongated aspect ratio (Fig. **1**A,B and Movie S1), and asymmetric closure dynamics, closing more rapidly from the rostral end compared to the caudal end (Fig. **1**A,C). We found that the cells surrounding the gap (row-1 cells) displayed robust elongation along the gap (Fig. **1**D), producing a predominantly rostro-caudal cellular orientation distribution (Fig. **1**E, blue). Interestingly, this orientation is lost in the next row of cells (Fig. **1**E, red).

We then proceeded to quantify different metrics of cell geometry, including cell area, aspect ratio, and the cell shape index (perimeter over the square root of the area [20–23]) (Fig. **1**F-H), for the first three cell rows using fixed mouse embryos ranging from the 13 to 17 somite stage. We found that row-1 cells have significantly smaller area than row-2 cells (18% smaller on average, Fig. **1**F). Additionally, we found that the mean cell aspect ratio (Fig. **1**G) and shape index (Fig. **1**H) decrease with the distance away from the gap. These measurements imply that the surface ectoderm robustly maintains highly elongated row-1 cells surrounding the gap and larger row-2 cells.

A key challenge in confluent epithelia is the visualization of individual cell shapes when surrounded by neighbors. To do this, we applied a mosaic, liphophilic dye which labels the entire cell membrane without crossing protein junctions between adjacent cells (Fig. **1**I). Visualizing the apical optical slices of mosaically labeled cells illustrates their polygonal shapes, equivalent to the shapes studied in simulations, and extension of membranous ruffles into the gap space directly ahead of row-1 cells (Fig. **1**J, * annotations). Mosaic labeling also visualizes broad protrusions which resemble cryptic lamellipodia, reaching beyond cells within the epithelium, towards the gap (Fig. **1**J-K). These protrusions, which to our knowledge have not previously been documented in this tissue, extend approximately 3-12 *µ*m ahead of row-2 and -3 cells towards the gap (Fig. **1**K-L), consistent with cryptic lamellipodia which power collective cell migration in other tissues [24]. Together, these analyses suggest that surface ectoderm cells around the HNP contribute to gap closure mechanics through actomyosin constriction and membrane protrusive activity, while maintaining robust patterning of cell morphology. Although we had previously modeled row-1 cells as the ones that actively crawl, our new data showing lamellipodium-like protrusions in cells away from the gap - together with the lack of established ECM ahead of row-1 - suggests row-2 and further cells are more likely to be the source of the crawling force.

## Cell crawling and purse-string tension are not sufficient to explain morphological patterning

To uncover the mechanisms regulating cell shape, cell orientation and collective motion during HNP closure in mouse embryos, we created an active vertex model for neuropore closure (Fig. **2**A). Specifically, we modeled the squamous surface ectoderm layer as a polygonal cell network in two dimensions, which achieves in-plane deformation in order to close the HNP. The mechanical energy of the surface ectoderm is given by [20, 25, 26]

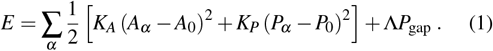

**Fig. 2.**
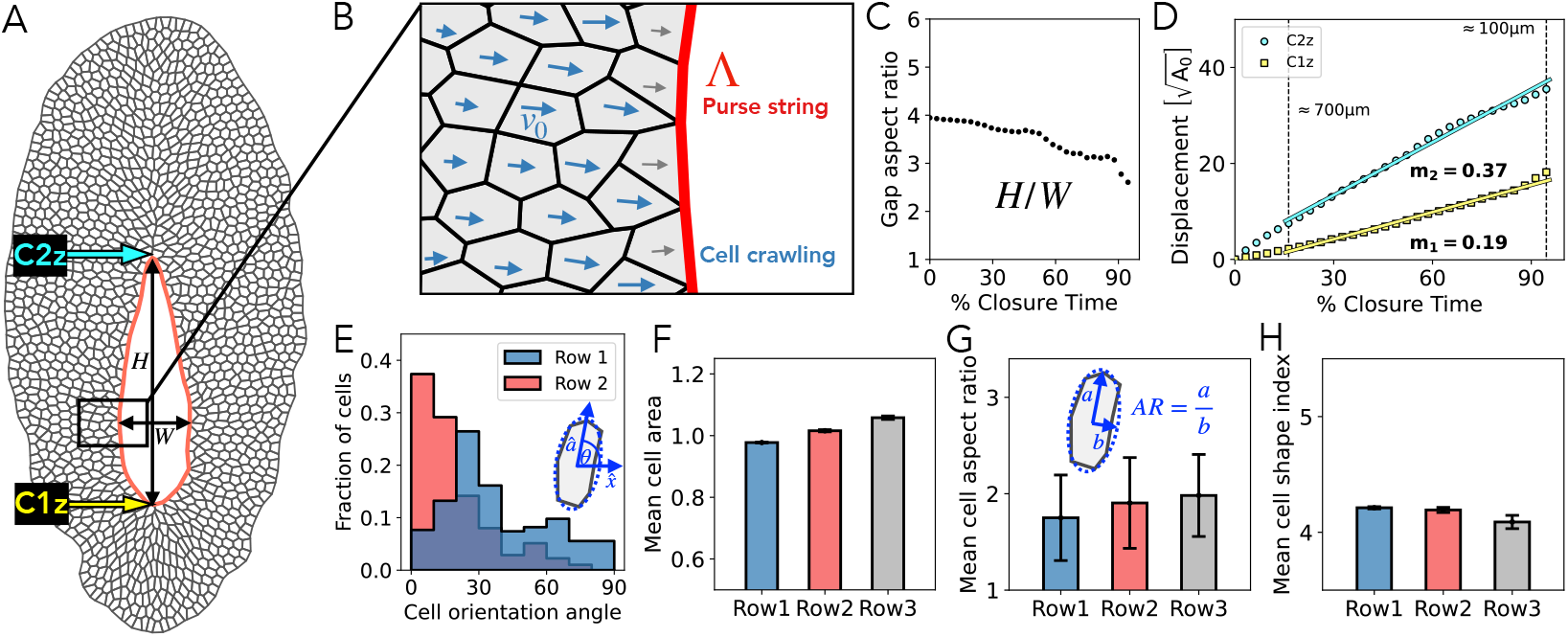
Cell crawling and purse-string contraction are not sufficient to explain cell shape anisotropy and patterning. (A) *In silico* tissue snapshot, representing a HNP of height *W* equal to 400 µm. The shape is evolved from an experimentally-determined starting point, with height *H*∼ 850 µm. C1z (yellow) and C2z (cyan) indicate the zippering points. The red contour highlights the high-tension purse-string along the HNP rim. (B) Schematic of the model for HNP gap closure. Active crawling of row-1 cells is restricted due to the paucity of extra-cellular matrix in the empty gap ahead of them. Second-row cells actively crawl towards the gap with a speed proportional to *v*_0_, guided by the normal-to-the-gap director of the first-row cells (illustrated by the black arrows). The active crawling speed of subsequent cells diminishes with increasing distance from the gap (illustrated by the length of the blue arrows). (C) HNP gap aspect ratio (*H/W*) versus percentage closure time. (D) Asymmetric closure of the zippering points C2z and C1z (cyan and yellow arrows in (A)) versus percentage closure time. Vertical black-dashed lines indicates time points at which the system represents a HNP whose height and width are approximately 700 µm and 100 µm, respectively. The solid cyan and yellow lines indicates the linear fit taking the times between the black-dashed lines. The cyan slope (*m*_2_) is approximately twice the yellow slope (*m*_1_). (E) Row-1 and Row-2 cellular orientation. Data account for information during the time window when row-1 consists of ∼ 13-36 cells, comparable to the analyses of embryos (Fig. **1**D,F-H). (F-H) Cell shape analysis considering the same time window as in (E). (F) Mean cell area per row. (G) Mean cell aspect ratio per row. (H) Mean cell shape index per row. The cell shape analysis in (E-H) excludes cells whose centers lie above C2z and below C1z. Error bars represent ±1 standard deviation.

The first two terms represent the elastic energy for each cell *α*, which penalizes changes in cell area *A*_*α*_ and perimeter *P*_*α*_, with respect to the target values *A*_0_ and *P*_0_. The area and perimeter elastic moduli are given by *K*_*A*_ and *K*_*P*_, respectively. The third term represents the purse-string mechanical energy, with a high tension Λ along the gap perimeter *P*_gap_ (red contour in Fig. **2**A,B). We modeled cell migration towards the gap as an active force acting on each vertex *i*, defined as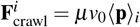 [27], where *µ* is the friction coefficient, *v*_0_ is the cell crawling speed, and ⟨**p**⟩ _*i*_ denotes the mean of the cell polarity vector **p**_*i*_ taken over the neighboring crawling cells. The position **r**_*i*_ of vertex *i* evolves in time as:

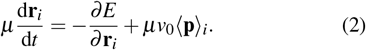

Due to the absence of an extracellular matrix ahead of the row-1 cells, it is assumed that row-1 cells cannot actively crawl into the fluid-filled void ahead of them, but assemble nascent matrix underneath as they displace towards the gap [6]. Row-2 cells actively crawl with a unit polarity vector pointing toward the gap. Finally, cells in the third row and beyond act as followers, with their polarity vectors **p**_*α*_ evolving in time through alignment with their immediate neighbors (denoted by the indices *β*) as:

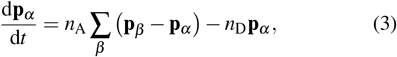

where *n*_A_ is the polarity alignment rate and *n*_D_ is the polarity decay rate [28]. These dynamics give rise to a polarity field oriented toward the gap, initiating in the second row of cells, and gradually decaying with distance (Fig. **2**B). This pattern is consistent with our previous visualization of collective displacement of cells in rows 1-3 [6].

Using the model described above, we simulated HNP closure using the parameter values given in Table I (see Supplementary Note 1, Fig. S1 and Movie S2). The model successfully captured the high aspect ratio of the gap throughout the closure process and the asymmetric closure rates, with the rostral zippering point C2z undergoing faster motion compared to the caudal zippering point C1z (Fig. **2**C,D). By varying *v*_0_ and Λ, we found that larger (smaller) values of *φ* = *µv*_0_*/*Λ led to a higher (smaller) gap aspect ratio (see Supplementary Note 1, Fig. S2). Notably, the gap-level features were also achieved when allowing row-1 cells to actively crawl, and in other variants of the model with different rates of polarity alignment and decay [6] (Supplementary Note 1, Fig. S2). This suggests that the gap-level dynamics are highly robust to model variations and depends primarily on the gap geometry.

**TABLE I.**
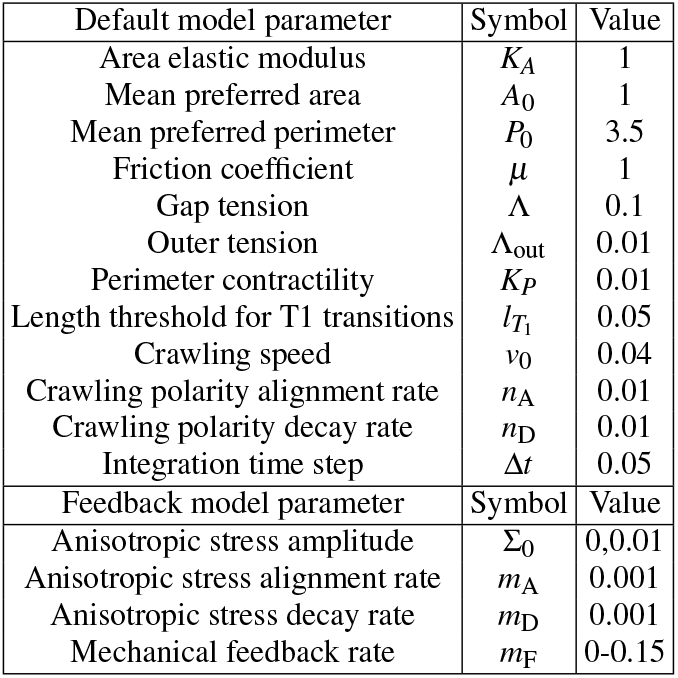
Default model parameters.

The vertex model, however, fell short of capturing the patterning of cell shape observed *in vivo*: it did not produce orientation or elongation of cells along the gap (Fig. **2**B,E,G,H and Movie S2). Cell shape index is similar in magnitude across different rows (Fig. **2**H), and much lower than experimentally measured values. Interestingly, the model was able to replicate other statistically significant experimental observations, such as row-2 cells have larger sizes than row-1 cells (Fig. **2**F) and larger perimeters than row-3 cells (Supplementary Note 3, Fig. S3), and cells positioned above C2z are larger than those below C1z (Supplementary Note 3, Fig. S3).

## Mechanical feedback model for anisotropic cell shape, motion and nematic organization

What causes the observed patterning of cell morphology? Elongation of row-1 cells, oriented along the gap (rostrocaudal axis), can mechanically originate from anisotropic stresses driving cell compression perpendicular to the gap (medio-lateral axis), coupled with extension parallel to the gap. Such anisotropic stresses could arise from acto-myosin meshworks generating active contractile or extensile stresses [29–31] We represent the average orientation of the actin fibers in each cell using the nematic order parameter **Q**_*α*_ (for cell *α*), which is a traceless and symmetric tensor that sets the direction of active stress **Σ**_*α*_ = Σ_0_**Q**_*α*_. Here Σ_0_ *>* 0 is a constant that sets the magnitude of contractile stress that the cell exerts on the surrounding environment. The work done by the anisotropic stress on the surrounding environment depends both on cell shape and nematic order [18], given by 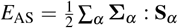, where **S**_*α*_ is the cell shape tensor (see Supplementary Note 2). This results in an active force exerted by the cells on the surrounding, 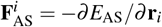, where **r**_*i*_ is the coordinate of the cell vertex *i*. By Newton’s third law, the force on the cell vertex is 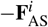. As a result, a cell will tend to elongate along the principal axis of **Q**_*α*_.

This raises the question of how the active stress orientation is regulated in the surface ectoderm. Orientational cues may originate from prepatterned gene expression, chemical signaling, or long-range external stresses resulting from the elongation of the body axis of the embryo. In Supplementary Note 4, we demonstrate how these models fall short in explaining the cell shape patterns around the HNP gap (see Figs. S9, S10). Alternatively, experimental evidence suggests that cells can polarize their actomyosin cytoskeleton in response to mechanical stresses [19, 32–36] This leads us to propose a mechanosensitive feedback model, in which the purse-string tension serves as a mechanical cue that drives cell shape changes by triggering a feedback between cellular shear stress ***σ***_*α*_ and nematic order **Q**_*α*_, such that **Q**_*α*_ aligns along the direction of maximum shear stress (Fig. **3**A). Specifically, **Q**_*α*_ is initially null and evolves in time through alignment with neighbors *β* (with a rate equal to *m*_A_), decay (with a rate equal to *m*_D_), and feedback with shear stress (with a rate equal to *m*_F_):

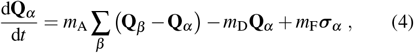

where all the rate constants are assumed to be positive.

**Fig. 3.**
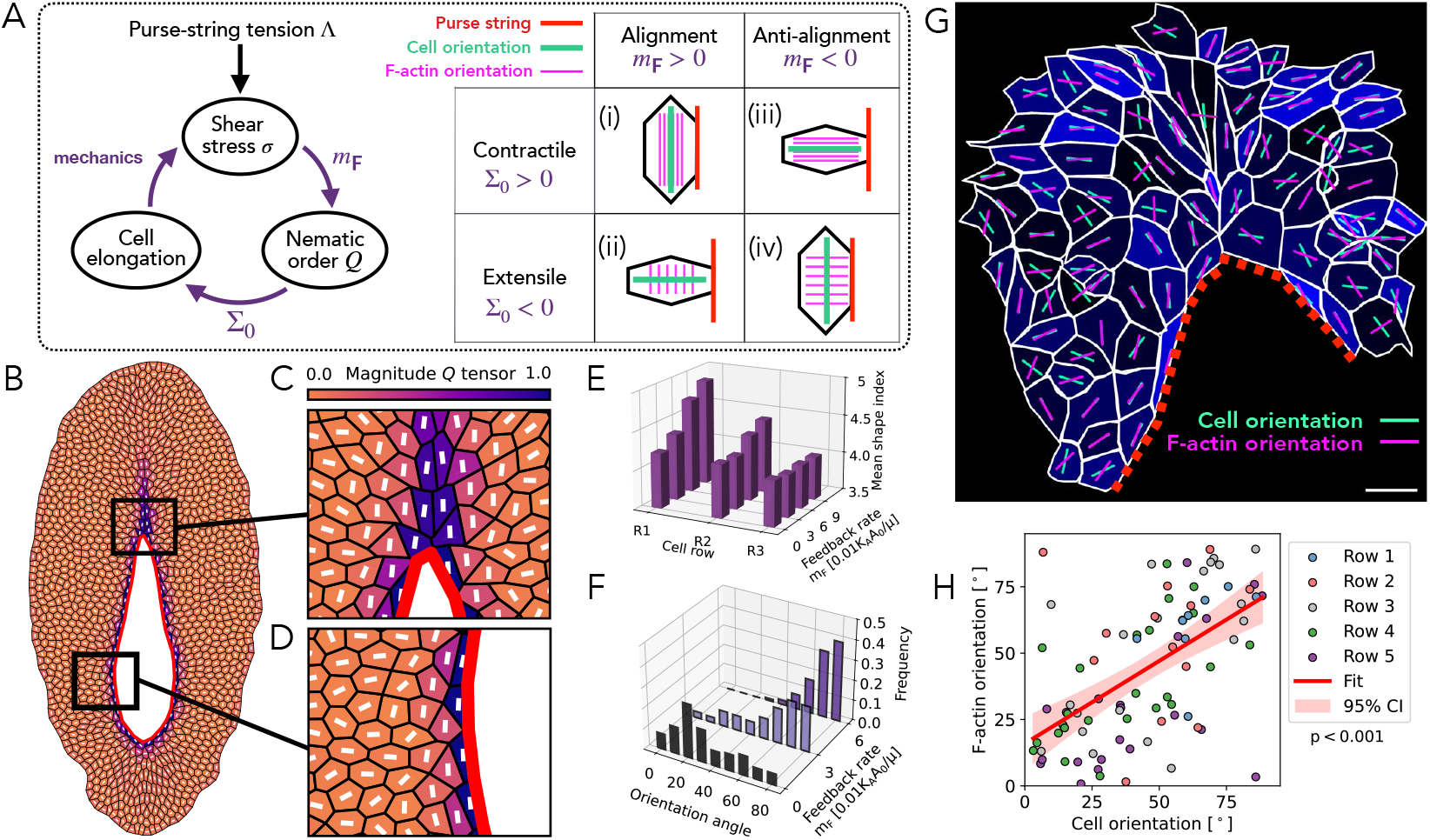
Mechanical feedback model for cell shape anisotropy and nematic patterning. (A) Schematic of the mechanical feedback model between cellular shear stress, nematic order **Q**, and cell elongation, controlled by the feedback rate *m*_F_ and the anisotropic stress magnitude Σ_0_. The purse-string tension Λ acts as a mechanical cue to trigger the feedback, driving cell elongation. (i-iv) Different combinations of row-1 cell orientations and nematic organization achieved by the model. (i) Contractile (Σ_0_ *>* 0) actin filaments aligned (*m*_F_ *>* 0) with shear stress; extensile (Σ_0_ *<* 0) actin filaments aligned (*m*_F_ *>* 0) with shear stress; (iii) contractile (Σ_0_ *>* 0) actin filaments anti-aligned (*m*_F_ *<* 0) with shear stress; and, (iv) extensile (Σ_0_ *<* 0) actin filaments anti-aligned (*m*_F_ *<* 0) with shear stress. (B-D) Application of the model to the HNP gap closure, with *m*_F_ = 0.08 and Σ_0_*/K*_*P*_ = 1. Tissue snapshot (B), top zoom-in (C), and lateral zoom-in (D) at 50% of closure time. The cell surface color represents the magnitude of the nematic order parameter **Q**, while the white bars indicate the nematic director. The red contour demarcates the HNP gap boundary. (E-F) Application of the model to the HNP gap closure with varying *m*_F_. (E) Mean cell shape index per row. (F) Row-1 cellular orientation distribution. Cell orientation angle is defined such that *θ* = 90^°^ indicates elongation along the rostro-caudal axis. The analysis conducted in panels (E-F) accounts for the time window when row-1 consists of ∼ 13-36 cells, comparable to the analyses of embryos and Fig. **1**C,D, and excludes cells above C2z or below C1z. (G) Illustration of surface ectoderm cell shapes and F-actin fibre orientation in cells around the Closure 2 zippering point. White outlines indicate the cell borders, green lines indicate the orientation of each cell’s long axis, magenta lines indicate the predominant orientation of their F-actin stress fibers, and blue shading indicates the coherency of their F-actin as defined by OrientationJ. See Supplementary Note 3, Fig. S4 for visualisation of the F-actin fibers. (H) Correlation of surface ectoderm cell long-axis and predominant F-actin stress fiber orientation calculated by OrientationJ (see Methods). Data points represent 88 cells in the embryo shown in Supplementary Note 3,Fig. S4 (representative of 4 equivalently-analyzed embryos).

The active force 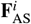 remains unchanged by a simultaneous reversal of the signs of Σ_0_ and *m*_F_. For example, when the active stress is extensile (Σ_0_ *<* 0) and anti-aligns with the shear stress 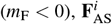 does not change. However, as we show later, experimental data do not support anti-alignment between nematic order and shear stress. Conversely, if Σ_0_ and *m*_F_ have opposite signs (Fig. **3**A.ii,iii), 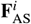 promotes cellular compression along the principal shear axis, inducing elongation perpendicular to the gap, which contradicts experimental observations.

The mechanosensitive feedback model successfully reproduced the elongation of row-1 cells along the gap (Fig. **3**B-D and Movie S3). We considered the limit of high nematic memory, *m*_A,D_ ≪ *m*_F_, based on the experimental observation of cells maintaining their elongated shapes (Fig. **1**A). The emergent pattern of the nematic field, representing the principal axis of **Σ**_*α*_, exhibits robust characteristics. First, in close proximity to the gap, lateral regions with low gap curvature exhibit a nematic field parallel to the gap (Fig. **3**B,D), inducing cell elongation along it. Second, once row-1 cells build their anisotropic stress, they are able to retain it for a long time after leaving the gap. This high nematic memory led to a stress field mostly perpendicular to the gap in the region above the zippering point C2z (narrow end, Fig. **3**B,C). In contrast, below the zippering point C1z (wider gap end), the nematic memory induced a transiently oblique nematic field Fig. **3**B. Over time, however, the nematic order parameter gradually aligned with the rostro-caudal axis (see Movie S3).

The model successfully reproduced the observed distribution of cell shape index per row (Fig. **3**E), and the rostrocaudal cell orientation in row-1 (Fig. **3**F) (see Table I for model parameters). Both row-1 cell elongation and rostrocaudal orientation increased with the feedback rate *m*_F_, and the magnitude of the active stress Σ_0_ (see Fig. **3**E,F, and Supplementary Note 3, Fig. S5). Additionally, the model predicted a more isotropic cellular orientation distribution in row-2 when increasing *m*_F_ and Σ_0_, in agreement with the experimental observations (see Fig. **1**E, and Supplementary Note 3, Fig. S6). Importantly, the mechanical feedback led to the expected cell geometrical patterns without interfering with the tissue-level dynamics (see Supplementary Note 3, Fig. S7).

To test the theoretical prediction of actin fiber alignment along the cell’s long axis (Fig. **3**B-D), we visualized F-actin distribution using high-resolution wholemount confocal microscopy of the surface ectoderm cells around the HNP in mouse embryos. Each cell has dense F-actin around its apical cell cortex (which largely overlaps with ZO1 staining and was excluded from analysis) and finer filaments resembling stress fibers basally (see Supplementary Note 3, Fig. S4). We quantified the coherence and average orientation of actin fibers and found a robust correlation between cells’ long axis and F-actin orientations (Fig. **3**G,H), which becomes particularly evident in cells with high F-actin coherence (see Supplementary Note 3, Fig. S4). These findings are consistent with our model assumption of a contractile nematic stress (Σ_0_ *>* 0, see Fig. **3**A). If instead Σ_0_ *<* 0, we would have expected actin fibers to be oriented perpendicular to cell’s long axis (Figs. **3**A), contrary to experimental observations.

## Tissue solidification maintains morphological patterns

The maintenance of cell patterns observable *in vivo* suggests suppression of cell rearrangements which would alter their positions over morphogenetically-relevant timeframes. Cell neighbor exchanges are directly linked to tissue material properties, with fluid tissues exhibiting more exchanges [20, 37]. We thus investigated how tissue material properties are regulated in the surface ectoderm to enable pattern maintenance.

Analysis of our simulations revealed an abundance of 4-fold vertices during HNP closure, mostly positioned around the gap and along the rostro-caudal midline region (Fig. **4**A). These 4-fold vertices represent stalled cell neighbor exchanges that arise during cell intercalations when the resolution of 4-fold vertices into tricellular vertices are energetically unfavorable (see Methods) [37]. Via live imaging we were able to identify the assembly and temporal stability of these structures above the rostral zippering point (Fig. **4**B and Movie S4), as predicted by our model (Fig. **4**C and Movie S5). Furthermore, we corroborated the robust presence of higher-order vertices across the surface ectoderm in fixed embryos (Fig. S10A).

**Fig. 4.**
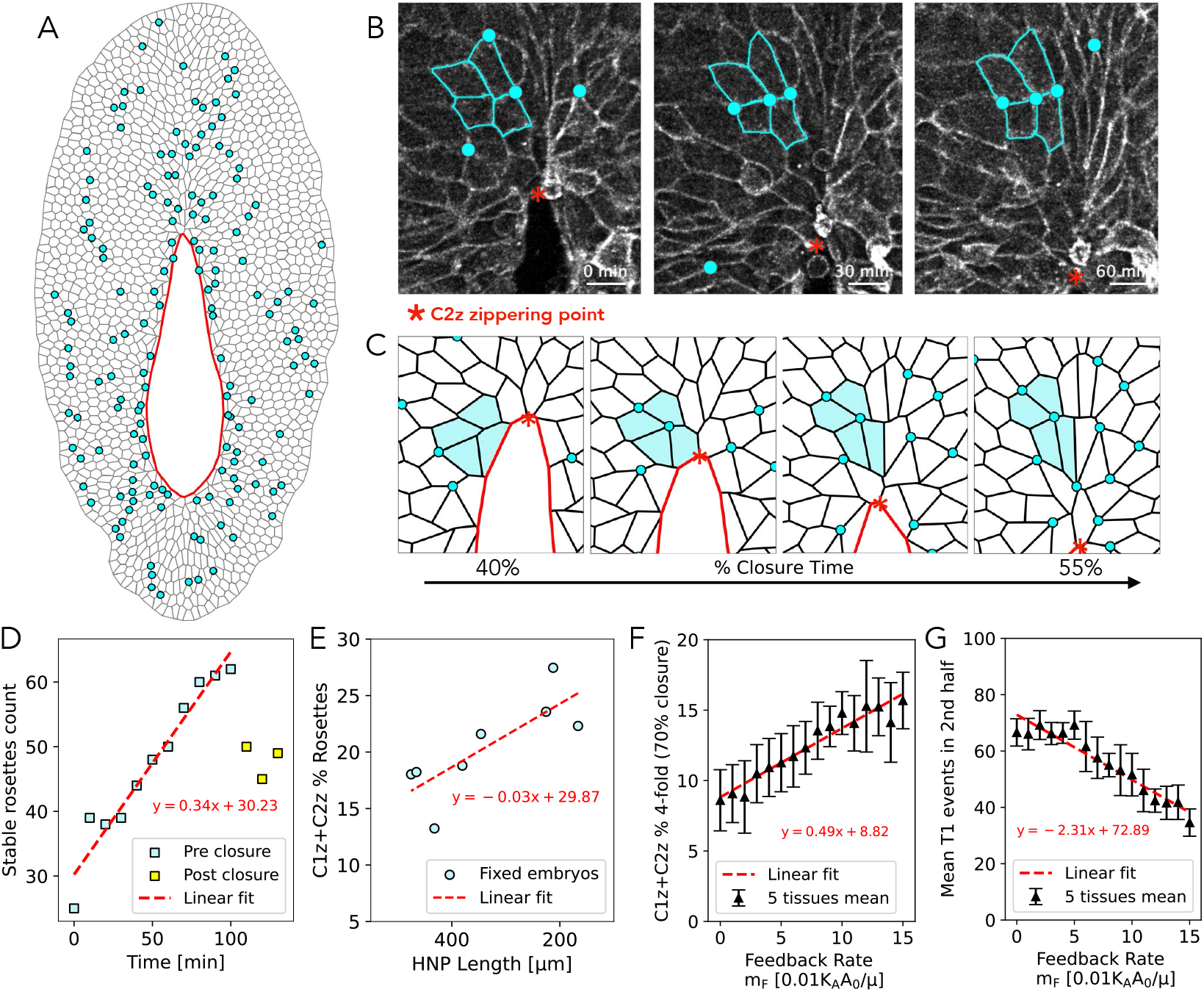
Mechanical feedback promotes tissue solidification by stalling cell neighbor exchanges. (A) Tissue snapshot using the mechanical feedback model to simulate HNP gap closure with *m*_F_ = 0.08 and Σ_0_*/K*_*P*_ = 1, at 50% closure. Cyan dots indicate 4-fold vertices present in the *in silico* surface ectoderm. (B) One-hour live imaging sequence of cells close to C2z, showing the presence of long-lived higher-order vertices (cyan dots). The four cyan cells indicate stable rosettes. Scale bar, 20 µm. (C) Simulation illustrating emergence of a stable 4-fold vertex equivalent to that live-imaging in (B). (D) Absolute count of stable rosettes over time during live imaging, in a region of size equal to 0.14 mm^2^. (E) Fraction of rosettes as a function of HNP length, defined as the number of higher-order cell junctions (≥ 4 cells) over the count of 3-fold vertices, found around C1z and C2z (covering about 225 vertices in each side) in 8 fixed embryos. (F-G) Tissue fluidity characterization in simulations with varying feedback rate. Each data point indicate the mean taken between 5 different simulations (varying initial conditions). Error bars represent ±1 standard deviation. (F) Fraction of rosettes at 70% of closure, calculated as in (E), considering circular regions centered at C1z and C2z with a radius equals 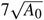. (G) Absolute T1 events during the second half of closure.

We found that, on average, the number of stable 4-fold vertices in the surface ectoderm increases over time (Fig. **4**D), in agreement with simulation results (Fig. S10B-C). Furthermore, we analyzed fixed embryos, finding a positive correlation between the fraction of rosettes around the zippering points and closure progression (Fig. **4**E). These results indicate that the surface ectoderm becomes more solid-like over time, as multicellular rosettes are signatures of tissue solidification [38].

To further characterize the effect of mechanical feedback on tissue solidity, we simulated closure with different initial configurations and quantified the fraction of 4-fold vertices around the zippering points at 70% of gap closure, as well as the absolute number of T1 events. Within our study range, the fraction of 4-fold vertices can increase by up to ∼ 100% (Fig. **4**F), while cell neighbor exchange events can drop by ∼ 50% (Fig. **4**G) for high values of *m*_F_, indicating that tissue solidity is enhanced by the feedback mechanism. Additionally, we found that HNP closure takes longer with increasing *m*_F_ (see Supplementary Note 3, Fig. S10E) - suggesting an evolutionary compromise between cell shape patterning and HNP closure speed.

## Anisotropic cell shape and motion patterns at the mid-line guides neuropore closure

Our mechanical feedback model identifies specific patterns of cell shape that can be directly tested with experimental data. Specifically, we found that the row-1 cells leaving the gap formed the midline region in later stages (Fig. **5**A), as seen in embryo live imaging (Fig. **1**A). These midline cells exhibit rostro-caudal elongation (Fig. **5**B). We found consistent results when simulating surface ectoderm midline cells (Fig. **5**C). Additionally, these results are consistent with experimental observations during spinal neural tube closure [39, 40].

**Fig. 5.**
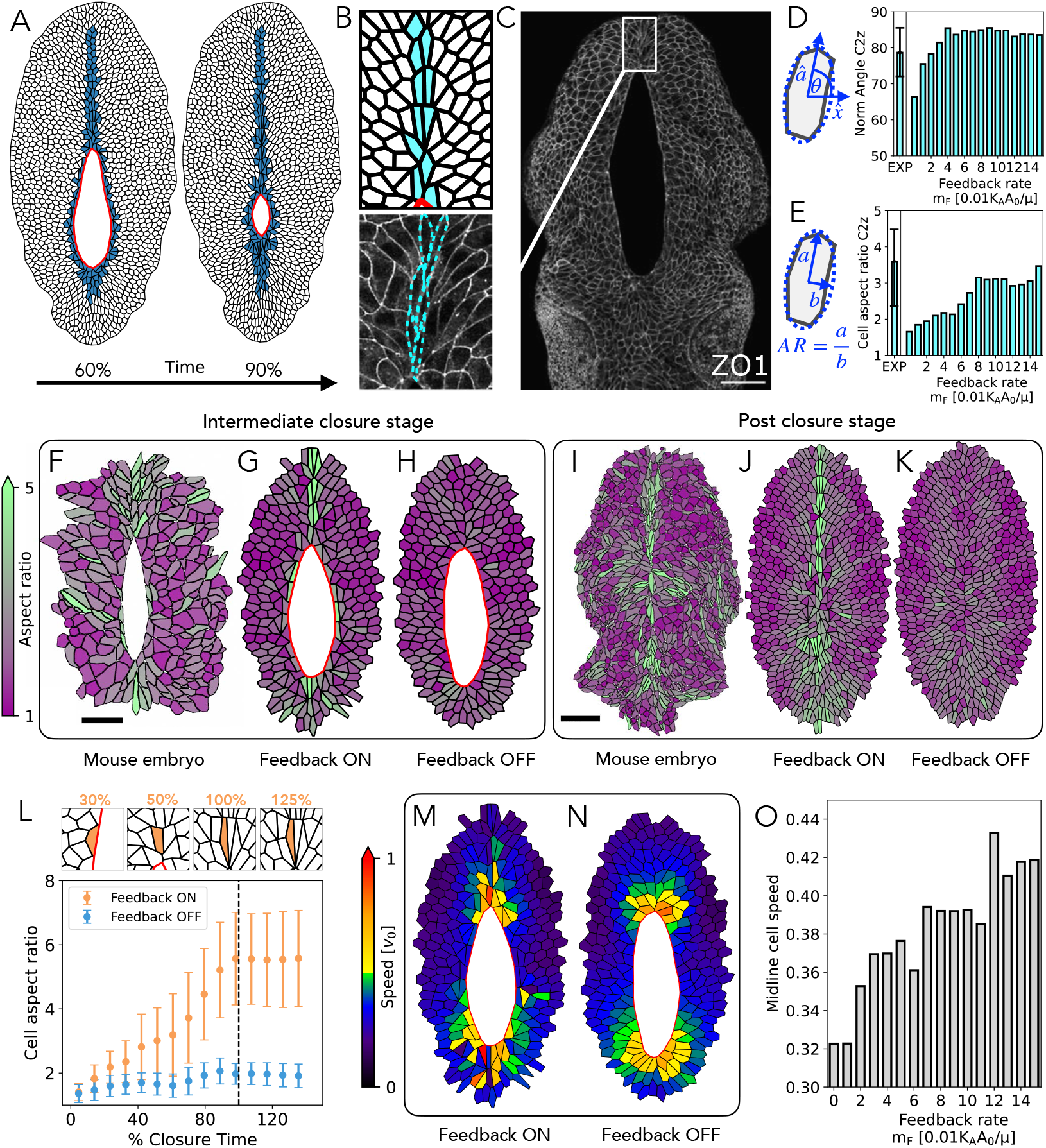
Mechanical feedback promotes midline cell elongation and rostro-caudal orientation. (A) Tissue snapshots using the mechanical feedback model to simulate HNP gap closure with *m*_F_ = 0.08 and Σ_0_*/K*_*P*_ = 1, at two distinct time points (60% and 90% closure). Blue cells represent the entirety of cells forming row 1 at *t* = 0. (B) Zoomed in snapshot of the midline cells situated just above C2z at 60% closure. Midline region is defined by a rectangle of height equals 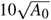 and width equals 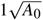, right above the zippering point. (C) HNP gap in a 14 somite fixed embryo, indicating midline cells as in the simulation shown in (B). Scale bar, 100 µm. (D-E) Comparison of the median rostral midline cell orientation and elongation as a function of the feedback rate *m*_F_, between experiments (50 individual cells taken from 8 fixed embryos at 15-17 somite stage), and simulations. Bars denote median value, while the error bars denote the 25/75 percentile range. Simulation data account for information during the time window when row 1 consists of 20 to 50 cells, mirroring experimental conditions. (F-H) Comparison of cell aspect ratio at an intermediate closure stage between a 15 somite mouse embryo (F), and simulations at 80% of closure, with (G) and without (H) the mechanical feedback. Scale bar, 50 µm. (I-K) Comparison of cell aspect ratio at full HNP closure. Scale bar, 100 µm. (L) Zoomed-in midline cell sequence at 4 different times. Comparison of the mean aspect ratio over time for cells that become midline cells (above and below the gap) between 30% and 90% of closure. Error bars represent ± 1 standard deviation. (M,N) Comparison of cell speed during HNP closure, with (M) and without (N) feedback. (O) Mean meadline cell (above and below the gap) speed for varying feedback rate.

We analyzed the morphologies of individual midline cells across 8 fixed embryos with open HNPs. To characterize the shape of these cells, we quantified cell orientation and aspect ratio. We found that the rostral midline cells were oriented along the rostro-caudal axis of the embryo (Fig. **5**D,E), reaching median aspect ratio values of 3.6. Additionally, our analysis of the caudal midline cells also revealed a rostrocaudal orientation, although with significantly lower aspect ratio (∼ 2.2) and greater orientation variation (see Supplementary Note 3, Fig. S8). We then performed HNP closure simulations and characterized the shape of midline cells. We found that the predicted midline cell shape and orientation quantitatively agreed with experimental data for sufficiently high feedback rate *m*_F_ (Fig. **5**D,E). Furthermore, the model with mechanical feedback reproduced the spatial pattern of cell aspect ratio observed in experiments, during and after closure (Fig. **5**F-K). In contrast, a weaker caudal midline cell elongation along with rostro-caudal orientation, emerge independently of the feedback mechanism (see Supplementary Note 3, Fig. S8).

Next we tracked all the prospective midline cells over time, and found that, on average, the mechanical feedback caused their aspect ratios to increase over time, and persist even after gap closure (Fig. **5**L), as corroborated in embryos (see Fig. **5**I). From a side-by-side comparison of cell aspect ratio and instantaneous speed computed from simulations (Fig. **5**G,M), we identified that the midline region is not only characterized by high cell aspect ratio but is also experiencing the largest instantaneous cell speed, with an average value that tends to increase with the feedback rate (Fig. **5**M-O). This underscores the fitness value of cell shape, such that high aspect ratio is associated with faster cell motion.

## Comparative morphogenesis reveals the functional role of purse-strings in cell patterning

Although neural tube closure is common to all vertebrates, different species may use alternative force-generating cell behaviors to achieve it. The rhombencephalic neuropore (RNP, Fig. **6**A) is the avian equivalent of the mammalian HNP. It closes through progressive medial apposition of two parallel tissue plates forming thin, slit-like gaps with often irregular “buttoning” contact points between them (Fig. **6**A) [41]. High resolution imaging of the RNP gap shows that it lacks F-actin purse-strings along the neural folds (Fig. **6**B), demon-strating an evolutionary difference from mice. We therefore hypothesized that the absence of mechanical feedback from contractile purse-strings would preclude gap-aligned elongation of leading edge cells observed in mouse embryos. Consistent with this, quantification of leading-edge cell orientation along the RNP gap reveals primarily gap-directed alignment (Fig. **6**C-D), in contrast to the rostrocaudal alignment observed around the mouse HNP, and closely resembling simulations of gap closure without purse-string (Fig. **2**E). Notably, the orientation of cells around the RNP is very similar to that predicted by the gap closure model without mechanical feedback. Simulating gap closure without high-tension purse-strings also produces corrugated leading edges (Fig. **6**E), reminiscent of the “buttoning” corrugations present during neural fold apposition in this species [42].

**Fig. 6.**
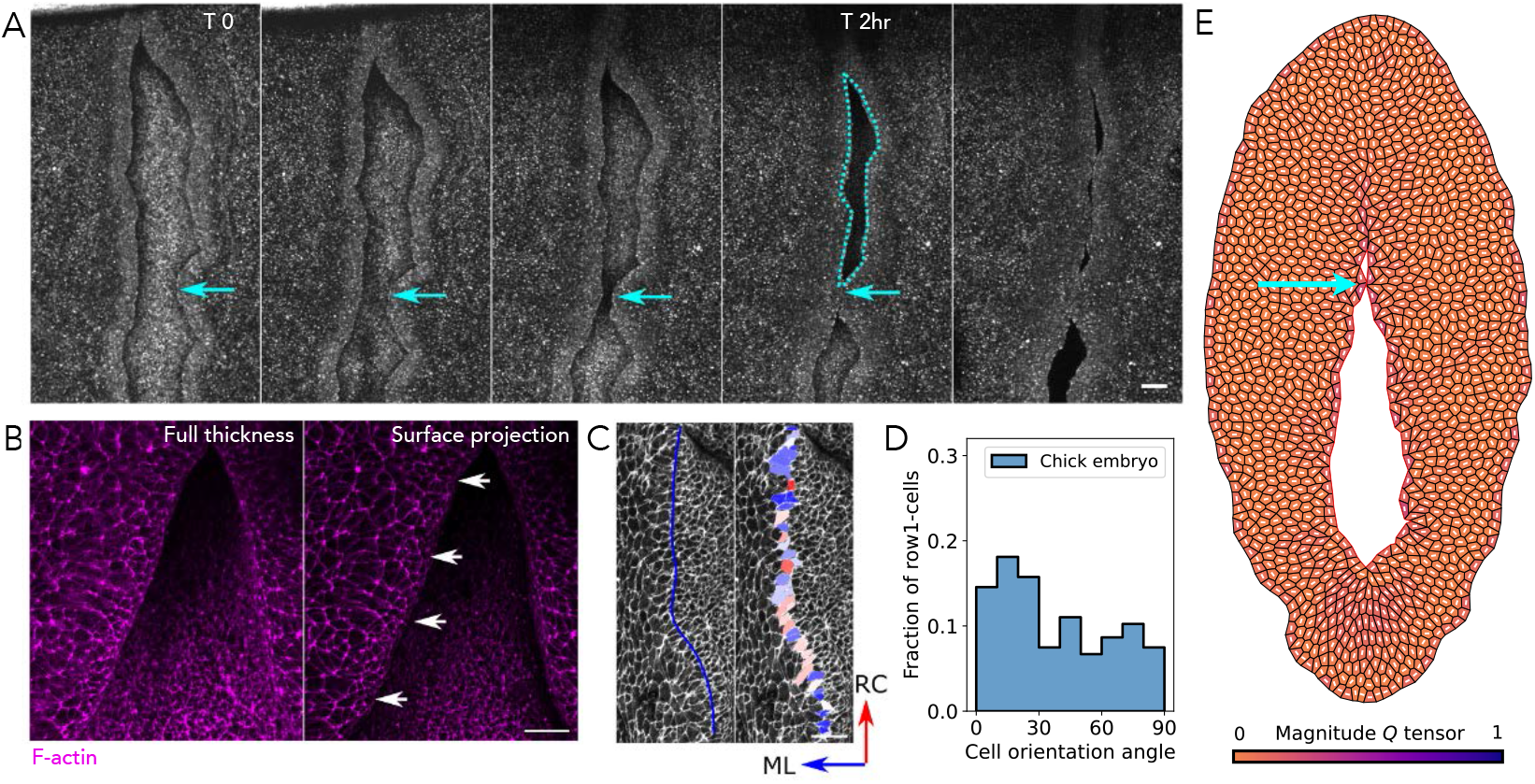
Evolutionary comparison: Chicken embryos do not use actomyosin purse-string to close their neuropores and their leading-edge cells are not elongated along the gap. (A) Reflection live imaging showing formation of the rhombocervical neuropore (RNP, cyan dashed line) through medial apposition of irregular neural folds (arrow). Note the slit-like gap shape. Scale bar, 50 µm. (B) Maximum intensity projection of a chick RNP showing the full tissue thickness, and a surface-subtracted projection to only show superficial staining. Note the absence of F-actin cables lining the neural folds (arrows). Scale bar, 25 µm. (C) Surface subtracted F-actin staining of an RNP neural fold. The blue line indicates the approximate demarcation between large, flat surface ectoderm cells on the dorsal tissue margin from small, apically-constricted neuroepithelial cells. Colour coding of the border cells indicates the orientation of the long axis (mediolateral ML, rostrocaudal RC). Scale bar, 25 µm. (D) Frequency distribution of the orientation of border cells (256 cells from 5 embryos). 0^°^ is mediolateral. (E) Tissue snapshot from a simulation of neuropore closure in a chicken embryo model lacking a supracellular purse-string. Gap tension is set to one-tenth of that used in the mouse HNP model, with mechanical feedback parameters *m*_F_ = 0.08 and Σ_0_*/K*_*P*_ = 1. The cell surface color represents the magnitude of the nematic order parameter **Q**, while the white bars indicate the nematic director. The thin red contour demarcates the HNP gap boundary. The cyan arrow indicates a gap-gap contact point occurring at a place different from the zippering points, as shown in (A). See Supplemental Information for specific modeling considerations for this simulation.

Thus, this “experiment of nature” demonstrates both the ability of embryos to close large tissue gaps without assembling supra-cellular purse-strings and that elongation of cells along the gap accompanies these high-tension structures rather than being a generalizable feature of neural tube closure.

## Discussion

Our work provides evidence that surface ectoderm cells exhibit predictable shape and nematic patterning during mouse HNP closure. We demonstrate that the observed gap dynamics and cell shape patterning can be explained purely by physical arguments. At the tissue-level, dynamics are robustly controlled by directed cell crawling and a purse-string mechanism. However, emergence and maintenance of cell morphological patterning requires active nematic stresses. We show that mechanosensitive feedback between cellular shear stress, nematic alignment, and cell shape can generate spatial patterns observed in experiments. Interestingly, the nematic feedback produces shape “memory” which extends the spatial and temporal outcomes of mechanical cues, in this case from the actomyosin purse-strings limited to the HNP rim. Inference of cellular properties from their shapes should therefore consider their mechanical histories. Predominantly parallel or perpendicular nematic patterning of leading-edge cells along closing gaps appears to be a generalizable property of many different systems (Fig. **7**).

**Fig. 7.**
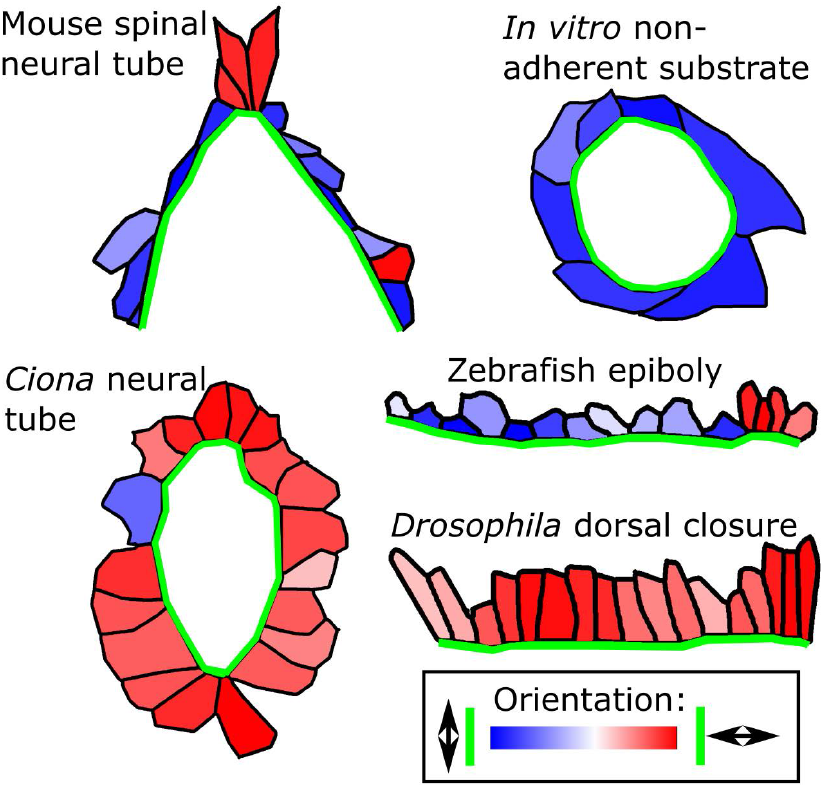
Schematic illustration of leading-edge cell orientation in various biological closure systems with different geometries and force-generating mechanisms. The mouse spinal neural tube zippering point has surface ectoderm cells aligned parallel to the gap edge, retaining that orientation as they leave the zippering point (26 somite embryo). MDCK cells closing over a non-adherent surface appear predominantly aligned along the gap (based on Nier et al. [46]). Zebrasfish epiboly also appears to include cells aligned along the gap (based on Bruce [49]). *Ciona* neural tube closure (based on Hashimoto et al. [50]) and *Drosophila* dorsal closure (based on De et al. [51]) appear to be two examples of gap closure with cells aligned perpendicular to the gap. Black outlines illustrate cell borders, green lines indicate the gap leading edge.

Ordered nematic patterning induced by the feedback mechanism leads to localized tissue solidification [22], which, while slowing HNP closure, actively preserves patterning within the surface ectoderm. It is not currently known whether this patterning guides development of derived structures, or is the result of evolutionary constraints imposing the requirement for mechanical feedback. Tissue solidification and deceleration of HNP gap closure is primarily driven by two factors: i) the movement of cells adjacent to the gap, whose dynamics is subject to a rim retraction force generated by anisotropic stress, and ii) the formation of persistent higher-order vertices. Consequently, gap closure times increase as feedback rate increases.

The mechanical feedback mechanisms cannot be neatly identified in embryos with available tools within the short period of HNP closure. Much of the actomyosin machinery is shared between cell types, such that pharmacological modulation dramatically changes tissue geometry [43, 44]. However, this mechanical feedback model is corroborated by our findings in chicken embryos, which lack the purse-string as the source of tension and do not show leading-edge cell elongation along the gap. The role of purse-string constriction has been formally tested in *Drosophila* embryos undergoing dorsal closure. High-tension purse-strings produce a straight leading edge which enables precise matching of the two halves of the embryo to fuse at a midline seam [45]. The chicken RNP, and *Drosophila* embryos lacking purse-strings, have less regular leading edges producing a wavier midline seam. A striking difference between these systems is that purse-string cells elongate perpendicular to the gap during *Drosophila* dorsal closure [45], as predicted from our models lacking mechanical feedback. Potential reasons for this difference include absence of feedback mechanisms in *Drosophila* or differences in their epidermal-amnioserosa interface versus the mouse surface ectoderm leading edge which extends mesenchymal-like protrusions into the gap ahead of them.

As such, mouse HNP closure is more akin to *in vitro* models of gap closure over non-adherent surfaces, which also appear to produce leading-edge cell elongation along the gap (Fig. **7**) [46, 47]. It is also likely that our findings in the HNP surface ectoderm apply in the spinal region of mouse embryos that also produces highly elongated midline cells [39]. Genetic disruption of potential mechanical feedback mechanisms involving myosin recruitment to cell-cell junctions abolishes nematic order of spinal midline cells [39]. In contrast, conditional surface ectoderm deletion of integrin *β* 1 produces larger cells which are rostrocaudally oriented with a high aspect ratio[48]. Thus, extrapolation between the cranial and spinal regions suggests that mechanical feedback leading to midline cell elongation may require cell-cell, more than cell-ECM interactions.

Overall, our study highlights crucial roles of mechanics as both a cue and driver of HNP closure. Future work could investigate the molecular mechanisms underpinning this feed-back and its generalizability to other biological systems, offering a bottom-up understanding of the physical principles governing epithelial patterning.

## Acknowledgements

SB acknowledges support from the National Institutes of Health (NIH R35 GM143042) and the National Science Foundation (NSF MCB-2203601). GLG acknowledges support from the Wellcome Trust (211112/Z/18/Z), Royal Society (RG \R2 \232082) and Leverhulme Trust (RPG-2024-147). EM acknowledges support from European Union’s Horizon 2021 Marie Sklodowska-Curie grant agreement no. 101067028. FPV acknowledges support from the NOMIS foundation. The surface subtraction macro is courtesy of Dr. Dale Moulding and available on GitHub (https://github.com/DaleMoulding/Fiji-Macros).

## Data availability

Source data are provided with this paper.

## Code availability

The computer code developed for this paper is available at: https://github.com/BanerjeeLab/HNP_model

## Methods

### Animal Procedures

Studies were performed under the regulation of the UK Animals (Scientific Procedures) Act 1986, the Medical Research Council’s Responsibility in the Use of Animals for Medical Research (1993), and were approved by UCL’s Animal Welfare Ethical Review Boards prior to issuing Home Office Licenses for the relevant work. C57BL/6J mice were bred in-house and used as plug stock from 8 weeks of age. mTmG mice maintained on a C57BL/6J background were as previously described [52] and tdTom fluorescence from homozygous mTmG embryos was used for live imaging. Mice were time-mated for a few hours during the day and the following midnight was considered E0.5. Pregnant females were sacrificed at E8.5. Embryos with 14-17 somites were analyzed.

### Live imaging, immunofluorescence, and image acquisition

Live imaging of tdTom-homozygous embryos was performed as previously described [6, 53]. Whole-mount immunostaining and imaging were also as previously described [54], using rabbit anti-zonula occludens (ZO)1 (Invitrogen 402200, 1:100) primary antibody. Secondary antibodies were used in 1:200 dilution and were Alexa Fluor-conjugated (Thermo Fisher Scientific). Alexa Fluor-647-conjugated Phalloidin was from Thermo Fisher Scientific (A121380). Images were captured on a Zeiss Examiner LSM 880 confocal using 10 x/NA 0.5 or 20 x/NA 1.0 Plan Apochromat dipping objectives and AiryFast. Images were processed with Zen 2.3 software and visualised as maximum projections in ImageJ/Fiji [55]. Linear adjustments were performed equally to all parts of each image. ImageJ/Fiji remove outliers processing was used to eliminate salt and pepper noise for segmentation.

### Image analysis

To visualise the surface ectoderm, the z-stacks were first surface-subtracted as previously described [56, 57] to only show the apical 2-5µm of tissue (macro courtesy of Dr. Dale Moulding available at https://github.com/DaleMoulding/Fiji-Macros). Tissue-level morphometric analysis was performed as previously described [6]. Cell morphometric analysis was performed using CellPose segmentation [58] executed in Napari, with manual correction, of ZO1 or phalloidin-stained surfaces. Standard cell shape parameters were measured using in-built functions in ImageJ/Fiji. Heatmap visualizations were created in ImageJ/Fiji, using cell outlines defined in the ROI manager to attribute cell shape properties to their greyscale intensity (scaled from 0-255).

F-actin orientation analysis was performed on surface-subtracted projections using OrientationJ [59–61]. Each cell was manually segmented within the cell cortex (eliminating the cortex from the analysis to avoid the longest cell borders biasing orientation quantification). F-actin filament staining intensity was equalized using the contrast-limited adaptive histogram enhancer (CLAHE) plugin (bin size = 50, histogram = 200, slope 3, slow processing) to account for variation in apparent intensity following surface-subtraction.

Surface projection was achieved using a previously-reported analysis pipeline [57] in which the top-most surface of the tissue is identified and a specified thickness of signal projected, excluding signal from cells below the surface ectoderm. Images were captured on a Zeiss LSM880 microscope with a 20x objective NA1 using AiryFast Super Resolution. Cell long axis orientation was calculated using the *fit ellipse* function in Fiji. F-actin orientation was calculated in individual cells using the plug-in OrientationJ [61], available online at https://bigwww.epfl.ch/demo/orientation/. F-actin stress fibres were accentuated before orientation analysis using the CLAHE function in Fiji (block size 50, bins 100, slope 3).

### Statistical Analysis

The statistical analysis shown in Fig. **1**F-H considers individual embryos as the unit of measure, and was performed in Excel 16.72. The rest of the statistical analysis considers individual cells as the unit of measure, and was performed in Python 3.9.7 (numpy 1.20.3). Comparison of two groups was by two-tailed Student’s t-test, paired by embryo where appropriate. Graphs were made in Python 3.9.7 (matplotlib 3.4.3) and are shown either as bar plots or as scatter plots. For bar plots, the bar shows the mean or median value, depending on the figure and described in each caption. The error bars indicate ± 1 standard deviation or the 25/75 percentile range, depending on the figure and described in each caption.

### Vertex Model Initialization

On average, the HNP forms at the 12-somite stage, exhibiting an asymmetric and elongated gap that is narrower at the rostral extreme (drop-like shape). The gap height and width reach up to 600 µm and 150 µm, respectively [6]. The closure of the HNP is typically completed by the 17-somite stage, taking 10 to 12 h. The typical cell area at early stages is ~ 200 µm^2^. To simulate HNP closure, we use an active vertex model where each cell is represented by a polygon whose vertices move in time following an overdamped equation of motion subjected to the forces of friction, tension, elasticity, and active crawling. We non-dimensionalized the equations using 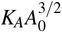 as the force scale, 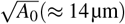 as the length scale, *µ/ K*_*A*_*A*_0_ as the time scale, and set *A*_0_ = 1, *K*_*A*_ = 1, and *µ* = 1.

The morphological cell patterning with row-1 cells exhibiting elongation along the gap is already observed at the 12-somite stage. Since in this work we aim at explaining the origin and maintenance of cell heterogeneity, we created an initial configuration that represent an earlier stage. Specifically, we generated a free (outer) boundary condition disordered tissue, composed of 1427 cells, with a gap that has a heightequals 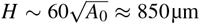, and a width equals 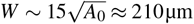.

### Numerical integration

The differential equations are integrated using the Euler integration method, with a time step fixed to ∆*t* = 0.05.

### Rules for T1 transitions

When a cell-cell junction connected by two 3-fold vertices becomes shorter than a threshold length 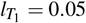, one of the vertices is removed, and the other is transformed into a 4-fold vertex, sustained by four shoulder junctions. At each time step, we attempt 4-fold vertex resolution in both the original (reversible T1-transition) and perpendicular (T1-transition) directions. The resolution is allowed only when the new configuration pushes the new 3-fold vertices apart in the direction of the largest repulsive force [37]. For the resolution attempts, we particularly consider the forces arising from elasticity and anisotropic stresses (both associated with energy terms), as well as the active crawling forces. Since the polarity field decays smoothly in the bulk region with cell-cell junctions, stalled 4-fold vertices indicate energetically unfavorable T1-transitions.

If a cell-void junction becomes shorter than a threshold length 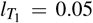, a T1-transition is performed instantaneously. Note that in this case, the junction is initially connected by two 2-fold vertices, whereas after the T1-transition, one of those vertices becomes a 3-fold.

### Mechanical Feedback Model

Our feedback model considers that cells are able to sense the local shear stress *σ*, and adjust the orientation of actin fibers in response. Here, we compute the tensor *σ* for a given cell *α* following the formalism previously described in Ref. [62],

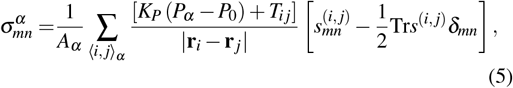

where the sum runs over adjacent vertices *i, j* belonging to the cell *α*, with positions **r**_*i*_ and **r** _*j*_, respectively, and *s*^(*i, j*)^ = (**r**_*i*_ − **r** _*j*_) ⊗ (**r**_*i*_ − **r** _*j*_). The variable *T*_*i j*_ can take the value *T*_*i j*_ = Λ if the junction is part of the gap boundary, *T*_*i j*_ = Λ_out_ if the junction is part of the outer boundary, or *T*_*i j*_ = 0, the junction is in the bulk. We neglect stresses arising from active crawling [63, 64] in our mechanical feedback.

The nematic orientation of F-actin within a cell *α*, **Q**_*α*_, leads to an anisotropic cellular stress given by **Σ**_*α*_ = Σ_0_**Q**_*α*_. Σ_0_ is a constant that sets the magnitude of the maximum contractile stress the cell exert over the substrate, and **Q**_*α*_ is a two-dimensional nematic tensor (traceless and symmetric tensor), defined by the magnitude *Q*_0*α*_ */*2 and the principal nematic director 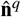. Then, the work performed by the anisotropic stress on the surrounding environment is given by 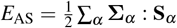, where **A** : **B** = *A*_*i j*_ *B*_*i j*_ = Tr (*AB)*^*T*^ (full tensor contraction), and **S**_*α*_ is the shape cell tensor given by 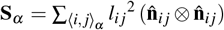, where **l**_*i j*_ = **r**_*i*_ − **r** _*j*_, *l*_*i j*_ = |**l**_*i j*_|, and 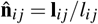. Then, the work can be written as the following:

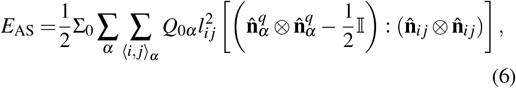

where 𝕀the two-dimensional identity matrix. By writing the previous tensors in the basis 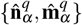, with 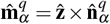, we obtain the following form for the energy cost:

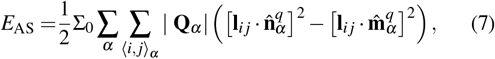

where the first sum is over the *α* cells to which the vertex *i* belongs to, and the second sum runs over the *j* adjacent vertices, belonging to the cell *α*. From the last quadratic-length form it is clear that the surrounding material will tend to elongate in the axis defined by 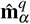 (extensile direction), with a magnitude of the stress given by Σ_0_ |**Q**_*α*_| = Σ_0_*Q*_0*α*_ */*2. Finally, each vertex *i* defining the cell boundary is subject to an extra force given by 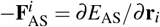.

