## Supplemental Information for "Anisotropic Cell Shape and Motion Coordinate Hindbrain Neuropore Morphogenesis"

### 1 Supplementary Note 1: Active Vertex Model Methods

#### 1.1 Choice of default model parameters

Laser ablation experiments [1] showed that recoil is approximately fivefold lower at surface ectoderm cell-cell junctions away from the gap, compared to cell junctions forming the gap boundary. This suggest that $\Lambda \sim 10K_P\Delta P$, with $\Delta P = P_{\text{bulk}} - P_0$. Additionally, we expect the outer boundary of the tissue to mechanically behave as the bulk of the tissue. Thus, we impose $\Lambda_{\text{out}} = K_P\Delta P = \Lambda/10$.

We find that cells far away from the gap tend to conserve their shape and position over time, which reflects a solid behavior (lateral cells in Fig. 1A). Additionally, data of 376 individual R3-cells taken from 9 different embryos, shows the 75-percentile shape index is given by $\sim 4.5$ (Fig. 2D). Thus, we decide to set $P_0 = 3.5$, and take $P_{\text{bulk}} \approx P_{\text{R3}}$, which yields $\Delta P = P_{\text{bulk}} - P_0 \approx 1$. Then, our conditions are simplified to $\Lambda = 10K_P$, and $K_P = \Lambda_{\text{out}}$. Thus, setting the value for $\Lambda$ sets it for $\{K_P, \Lambda_{\text{out}}\}$. We additionally set the ratio $\mu v_0/\Lambda = 0.4$, which we find to be good to describe HNP dynamics. Then, we simulated HNP closure for several values of $\Lambda$, subject to all the previous conditions for $\{K_P, \Lambda_{\text{out}}, v_0\}$. This procedure allow us to play with the ratio between cell elasticity and the rest of the forces acting over the vertices. We try different regimes and compare the outcomes to robust gap level features observed in fixed and live-imaged mouse embryos (Fig. S1). These include the maintenance of a high gap aspect ratio throughout closure (Fig. S1A), and the asymmetric closure dynamics with the caudal zippering point moving faster (Fig. S1B). Firstly, we find that the gap aspect ratio decreases for increasing $\Lambda$, getting more circular-like that what was observed in the live-imaged embryos (Fig. S1C). Secondly, the difference in net displacement of the zippering points increases with $\Lambda$ (Fig. S1D). To capture both experimental robust gap-level observations, we decide to set $\Lambda = 0.1$ for all the simulations.

The cell crawling dynamics is chosen such that it creates a polarity field pointing toward the gap. As explained in the main text, row-1 cells do not actively crawl. Nevertheless, these cells play a pivotal role in directed cell migration through their interaction with the cells of the subsequent row. Specifically, the polarity vector of a row-2 cell has unit magnitude, and its direction is given by the mean normal to the gap (inward) $\hat{\mathbf{n}}_{\text{R1}}$ of its row-1 cell neighbors. As a result of cell polarity-polarity alignment (with rate $n_{\text{A}}$) and decay (with rate $n_{\text{D}}$), the polarity field starts in the second row of cells, and decay with the distance from the gap. To set the default values we first simplify the system. If we think in a straight gap boundary front, the dynamics described for the polarity field can be translated to a one-dimensional equation given by $\partial_t p(x,t) = n_{\text{A}}\partial_x^2 p(x,t) - n_{\text{D}}p(x,t)$. We can assume that the

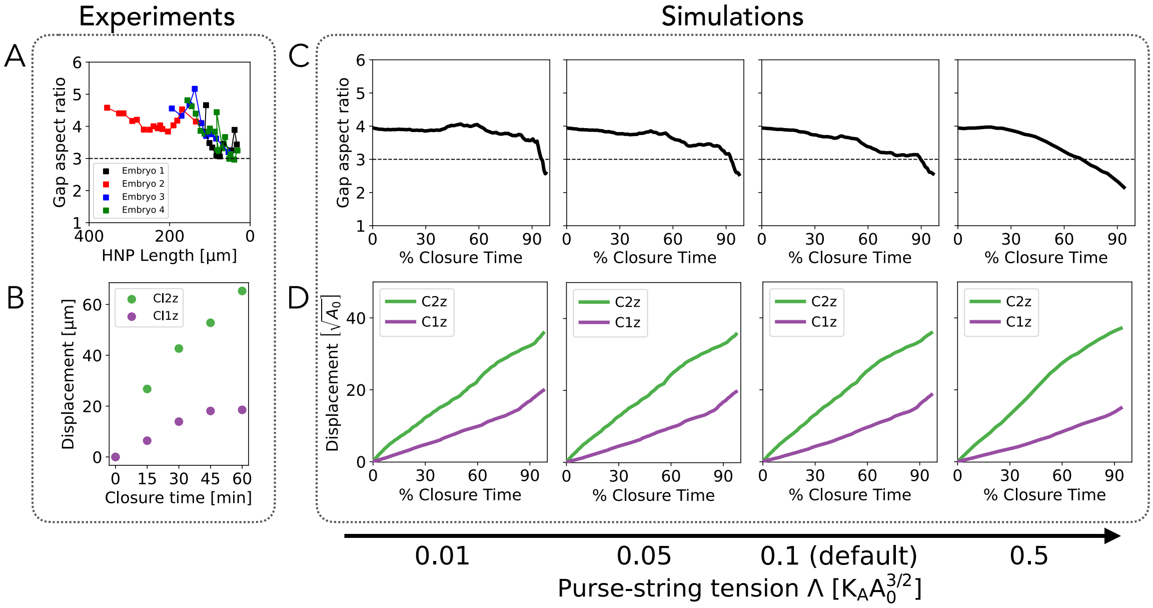


Fig. S1: **Gap-level dynamics during HNP closure.** (A,B) Experimental data derived from observations on live-imaged mouse embryos. (A) Gap aspect ratio versus HNP length, $n = 4$. Embryos 1-3 were previously published [1], and Embryo 4 is shown in Fig. 1A. (B) Quantification of C2z and C1z (zippering points) displacement over time (from $T = 0$, Fig1.A). (C,D) Simulation results for varying values of the purse-string tension, with the system subject to low cell junctional tension ($K_P = \Lambda_{\text{out}} = \Lambda/10$), and $\mu v_0 = 0.4\Lambda$. (C) Gap aspect ration versus percentage closure time. (D) Asymmetric C2z and C1z displacement versus percentage closure time.

front is defined by the position of the second row, and therefore $p(x = \text{front}, t) = 1$. From this one-dimensional perspective we can easily see that the values of the rates $n_{\text{A,D}}$ will set the time it takes the polarity field to be built. However, for times much longer than $1/n_{\text{A,D}}$ the polarity field reach $\partial_t p(x,t) \approx 0$. The steady-state solution of the polarity field is given by $p(x) = \exp\left(-\sqrt{n_\text{D}/n_\text{A}}x\right)$, where $x$ is the distance from the front. Thus, $\sim 3\sqrt{n_\text{A}/\left(n_\text{D}A_0\right)}$ gives an estimate of the number of rows that will be actively crawling toward the gap.

In our numerical simulations, we impose the following assumptions: i) the evolution of the cell polarity is slower than the motion of a row-2 cell in total isolation (due to active crawling only); ii) only a couple of rows are actively crawling during HNP closure. Both conditions are satisfied by considering $v_0 > n_{\text{A,D}}$, and $n_\text{A} = n_\text{D} = 0.01$.

#### 1.2 Gap shape control by varying $v_0$ and $\Lambda$

We test varying values of the cell speed factor $v_0$ and the purse-string tension $\Lambda$, showing both mechanisms control the gap aspect ratio in time (Fig. S2A-C). Specifically, the gap becomes more circular-like for increasing (decreasing) $\Lambda$ ($v_0$). We additionally show the gap shape control with a model in which the first row can actively crawl and act as the leader of polarity (Fig. S2D-F).

#### 1.3 Additional gap-junction adhesion consideration for the low gap-tension simulation

To mimic the gap closure that occurs in chick, we reduce the gap tension to 1/10 of the HNP *in silico* value. This causes consecutive large gap junctions to come very close facing each other with a small angle in between (mostly at the zippering point). In these cases, we allow fast adhesion of those junctions if the angle is less than 10°, resembling a buttoning closure mechanism. If the gap junctions that come together are not consecutive, then we stop the simulation before the vertices undergo cell crossing.

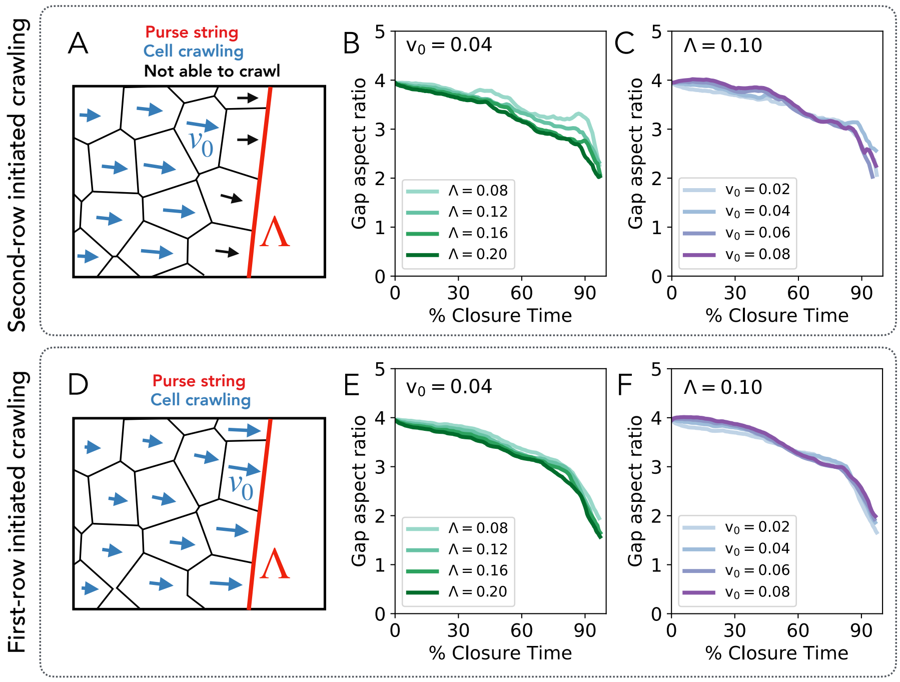


Fig. S2: **Purse-string tension and active crawling regulate gap shape in a second and a first-row initiated crawling model.** (A-C) Second-row initiated crawling model. (A) Model schematic. (B) Gap aspect ratio against percentage closure time, for different gap tension with fixed crawling speed. (C) Gap aspect ratio against percentage closure time, for different cell crawl speeds with fixed gap tension (D-F) First-row initiated crawling model. Same representation as (A-C).

### 2 Supplementary Note 2: Mechanical feedback model

#### 2.1 Choice of mechanical feedback model parameters

In our mechanical feedback framework, cells are able to build up a nematic tensor $\mathbf{Q}$ that generates an anisotropic medial stress $\mathbf{\Sigma} = \Sigma_0 \mathbf{Q}$. $\mathbf{Q}$ changes in time via mechanical feedback (with a rate $m_{\mathrm{F}}$), neighbor alignment (with a rate $m_{\mathrm{A}}$), and decay (with a rate $m_{\mathrm{D}}$). For the model, we initially set the nematic order parameter to be zero everywhere, and additionally consider that there is a maximum anisotropic medial stress that cells can achieve. We implement this idea by setting the maximum magnitude of $\mathbf{Q}$ to one. Therefore, the maximum magnitude of $\mathbf{\Sigma}$ is given by $\Sigma_0$. We take $\Sigma_0$ to be similar to $K_P$, which is the other cell level quantity with the same units. Thus, we test $\Sigma_0/K_P = \{0.5, 1\}$.

The experimental observation of non row-1 cells conserving their elongated shape and orientation while moving over time (Fig. 1A) motivates us to work in a regime of i) high nematic memory, $m_{\mathrm{A}} = m_{\mathrm{D}} = 0.001$, and ii) fast build up of $\mathbf{Q}_\alpha$ (compared to the crawling polarity field dynamics). Particularly, we use $m_{\mathrm{F}}$ ranging from 0 to 0.15, in multiples of $\Delta m_{\mathrm{F}} = 0.01$.

#### 3 Supplementary Note 3: Additional experimental analysis and results of the theoretical model

**Cell-level heterogeneity within the surface ectoderm in embryos and *in silico* model.** Figure S3 shows the cell-level characterization of different groups of cells. The analysis includes cell area, perimeter, and shape index. While in the absence of the feedback mechanism our model is able to reproduce statistically significant trends observed in fixed embryos, the feedback is required to reproduce the mean cell perimeter and shape index decreasing with the distance from the gap.

**Surface ectoderm cell and F-actin orientation analysis.** Figure S4 shows an example of input (surface-subtracted projection) and output of the orientation analysis using OrientationJ. We find a negative correlation between the coherence of F-actin within a given cell, and the discrepance between cell and F-actin alignment (calculated as cell orientation angle minus F-actin orientation angle).

**Cell elongation magnitude per row depends on the value of anisotropic stress amplitude $\Sigma_0$.** Figure S5 shows the mean shape index for rows 1, 2, and 3, as well as the row-1 cellular orientation distribution, for varying values of feedback rate and $\Sigma_0/K_P = 0.5$ (50% of the value used in the main text). While this low value of anisotropic stress is able to achieve rostro-caudal orientation of row-1 cells, it is not enough to drive the expected elongation per row.

**Feedback model captures local rostro-caudal elongation.** Figure S6 shows that for both high feedback rate and anisotropic stress amplitude, a rostro-caudal orientation is achieved in row-1 which is lost in the subsequent row of cells around the gap.

**Effect of feedback mechanism on gap-level dynamics during HNP closure.** Figure S7 shows that increasing the feedback rate does not qualitatively affect the high aspect ratio of the HNP during closure or the asymmetric progression (faster on the rostral end). However, as the feedback rate $m_{\mathrm{F}}$ increases, we observe a tendency for the gap to become more circular.

**Caudal midline cells morphology is feedback-independent.** Figure S8 shows that the experimental rostro-caudal elongation of caudal midline cells is reproduced in the absence of the feedback mechanism. Increasing the feedback rate $m_{\mathrm{F}}$ has minimal impact on the overall geometric outcome

**Comparison of midline cell morphodynamics using different nematic models.** Figure S9 compares the average midline cell morphology over time across 3 biological-motivated nematic models. Results indicate that a time-dependent nematic field is necessary to reproduce experimental cell morphologies. The feedback model (shear stress-dependency), unlike the nematic pre-patterned II model, predicts an increasing mean cell aspect ratio during HNP closure.

**Effect of feedback on tissue solidity during *in silico* HNP closure.** Figure S10 presents three indicators of tissue solidification during gap closure: i) slower T1 event rate, ii) an increased rate of stalled 4-fold vertex formation, and iii) a longer time to complete gap closure.

#### 4 Supplementary Note 4: Discussion of alternative models

Alternative biophysical origins for the cell shape patterning observed in the surface ectoderm during HNP closure are also possible. Here, we compare different hypotheses based on the morphodynamics of midline cells. One hypothesis is that this organization is influenced by the long-range stresses caused by the elongation of the embryo's body axis. However, our experiments show that the preferential rostro-caudal cell orientation is lost in the row-2 cells surrounding the gap. Since the elongation of cells along the gap is lost by the second row of cells, a global mechanism, such as tension generated by body axis elongation, is unlikely to be sufficient to explain this localized patterning of cell shape.

However, the localized cell shape and orientation organization could be genetically regulated during development. To test this idea, we perfomed *in silico* simulations where cells begin with isotropic shapes, but the nematic director of row-1 cells is pre-patterned to align with the gap boundary (Fig. S9). We considered two scenarios: one where the nematic director remains fixed throughout closure (infinite nematic memory limit) and another where cells can modify their intracellular organization through alignment and decay. Both models reproduce the rostro-caudal orientation of midline cells within the experimental range (Fig. S9H-J,L), but they differ in terms of elongation magnitude and dynamics (Fig. S9K-N). In the fixed pre-patterned nematic model, the predicted caudal midline cell elongation is almost twice the experimentally observed values in mouse embryos, with mean elongation increasing during closure but decaying post-closure (Fig. S9N). In contrast, in the dynamic pre-patterned nematic model, while midline cell elongation lies within the experimental range, these cells achieve maximum elongation at approximately 40% gap closure, after which elongation starts to decreases over time. These results indicate that a mechanically dependent nematic evolution, as achieved through our mechanical feedback model, is necessary to reproduce the experimental observations.

Additionally, we tested an alternative model lacking the feedback mechanism, in which the initial condition represents a system with 50 row-1 cells exhibiting elongation along the gap (Fig. S11A–C), mimicking the geometry observed in the earliest embryos analyzed here. We found that, while the gap morphodynamics remained largely unchanged compared to those produced by our mechanical feedback model (Fig. S11D), the cell-level rostro-caudal orientation and elongation (Fig. S11E–G) weakened over time. Moreover, although the midline cells (Fig. S11H) maintained their rostro-caudal orientation (Fig. S11I), their elongation was significantly reduced above the C2z closing point (Fig. S11J), reaching approximately $\sim 50\%$ of the experimentally observed value (Fig. S9K), and displaying an opposite relationship between cell aspect ratio above and below the gap (Fig. S9K,M; Fig. S11J).

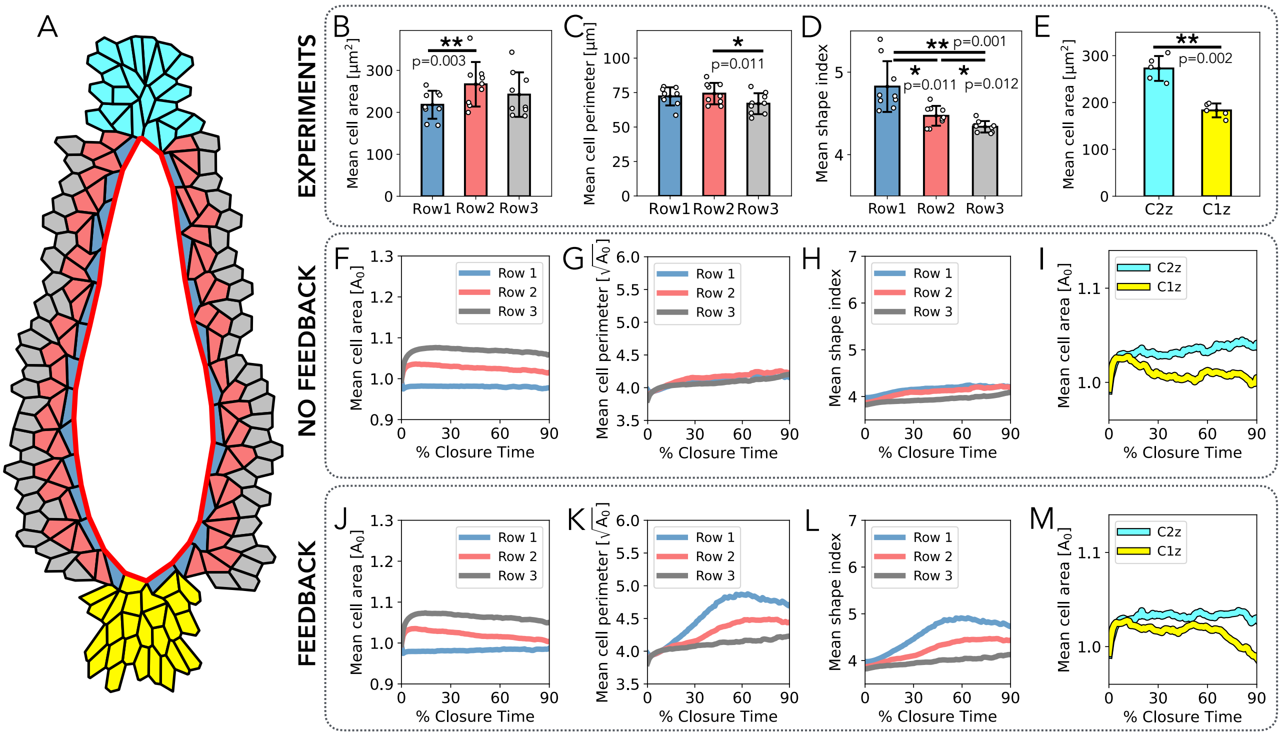


Fig. S3: ***In silico* model reproduces cell-level heterogeneity observed in the surface ectoderm cells.** (A) Simulated tissue snapshot showing the group of cells analyzed in experiments and simulations. Row-1 (blue), row-2 (red), and row-3 (gray) groups neglect the cells whose centers lie above or below the gap. C2z and C1z groups represent the cells whose centers lie in a squared region of $5 \times 5\sqrt{A_0}$, right above and below the gap, respectively. (B-E) Experimental data derived from observations on fixed mouse embryos. Mean cell area (B), perimeter (C), and shape index (perimeter over the squared root of the area) (D) for the first three rows, $n = 9$. (E) Mean cell area for the C2z and C1z groups of cells (green and purple, in A), $n = 5$. Error bars indicate ± standard deviation. The statistical analysis shown in (B-C) considers individual embryos as the unit of measure. Individual data points are represented with white dots.$*P < 0.05, **P < 0.01$. (F-I) Simulation results without feedback. Mean cell area (F), perimeter (G), and shape index (perimeter over the squared root of the area) (H) for the first three rows, versus percentage of closure time. (I) Mean cell area for the C2z and C1z groups of cells (green and purple, in A). (J-M) Simulation results with feedback, with $\Sigma_0/K_P = 1$ and $m_{\mathrm{F}} = 0.08$. Same representation as (F-I).

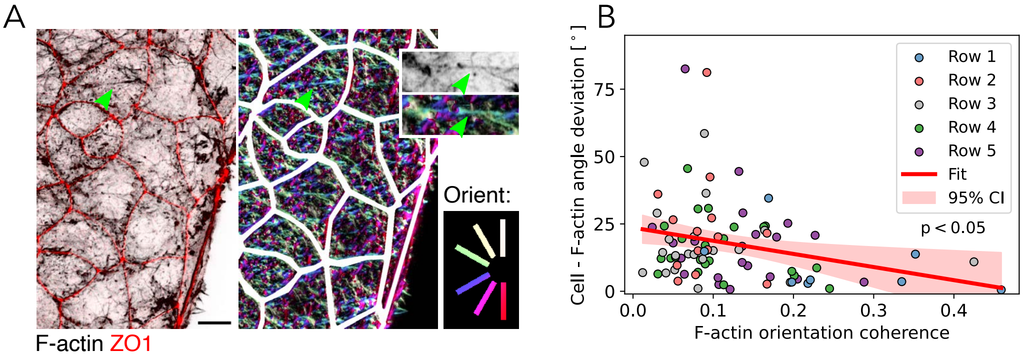


Fig. S4: **Surface ectoderm cell and F-actin orientation measurement.** (A) Left: Surface-projected [2] image showing ZO1 in the apical 2 µm and F-actin in the apical 5 µm of the surface ectoderm cell layer at the C2z of an embryo with 15 somites. Scale bar, 20 µm. Right: F-actin orientation obtained using the plug-in OrientationJ [3]. (C) Negative correlation between cell - F-actin angle deviation and F-actin orientation coherence. Data points represent 88 cells in the embryo shown in Fig. 2F.

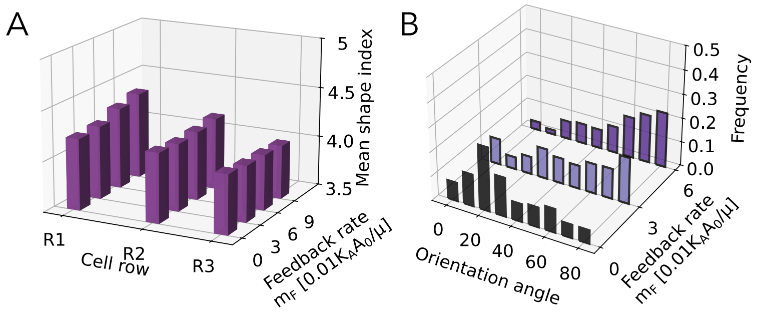


Fig. S5: **Application of the model to the HNP gap closure, with $\Sigma_0/K_P = 0.5$, and varying $m_\mathbf{F}$.** (A) Mean shape index per row. (B) Row-1 cellular orientation distributions. Analysis accounts for the same time window as Fig.1C,D, and excludes cells above C2z and below C1z.

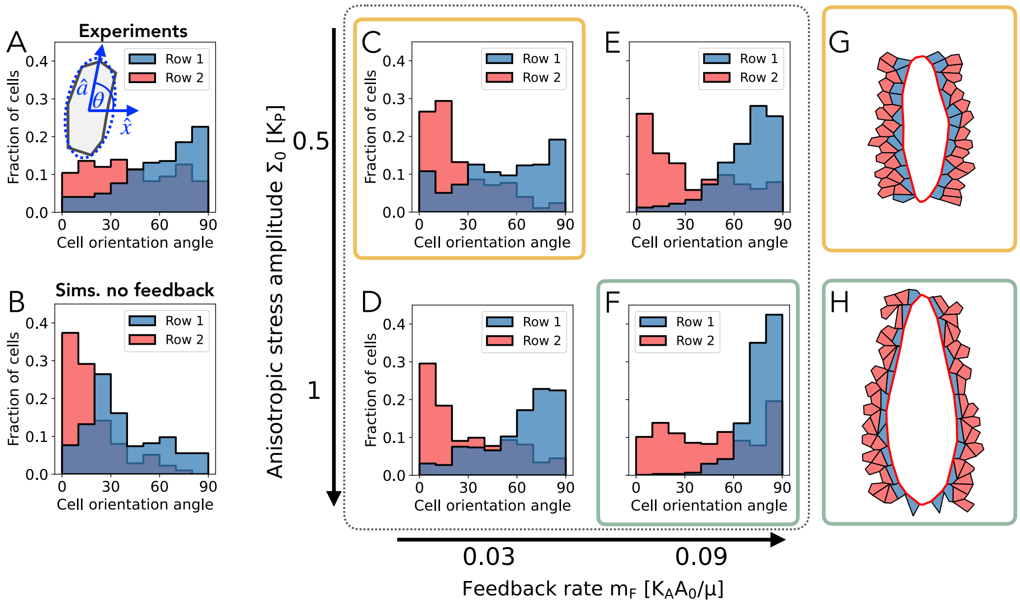


Fig. S6: **Feedback mechanism qualitatively reproduces experimental row-1 and row-2 cellular orientation distributions.** (A) Row-1 and row-2 cellular orientation distributions on fixed moused embryos. The normal vector $\hat{\mathbf{a}}$ defined the direction of cell orientation. Orientation angle is defined such that $\theta = 90°$ indicates cellular elongation along the rostro-caudal axis. The analysis considers 221 individual cells, neglecting cells above C2z and below C1z, from 9 fixed embryos. (B) Row-1 and Row-2 cellular orientation obtained in the simulation without feedback. Simulation data accounts for information during the time window when row 1 consists of 20 to 50 cells, mirroring experimental conditions. (C-F) Row-1 and Row-2 cellular orientation distribution for varying feedback rate $m_{\mathrm{F}}$ and anisotropic stress amplitude $\Sigma_0$. Analysis accounts for the same window of time as B, and excludes cells above C2z and below C1z. (G,H) Simulated tissue snapshots showing row-1 (blue) and row-2 (red) cells at 50% of the time window considered in the simulation represented by (C) and (F), respectively.

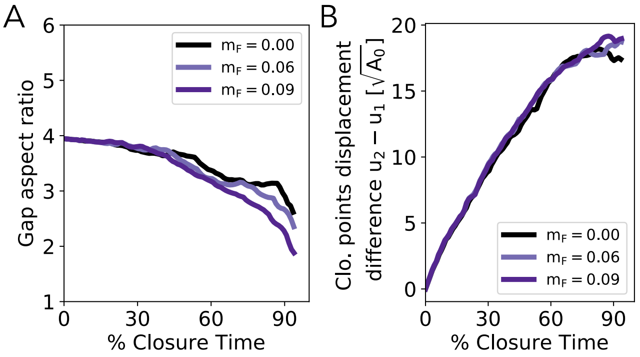


Fig. S7: **Feedback mechanism minimally alter gap level dynamics during HNP closure.** Application of the model to the HNP gap closure, with $\Sigma_0/K_P = 1$, and varying $m_{\mathrm{F}}$, in units of $0.04 K_A A_0/\mu$. (A) HNP gap aspect ratio versus percentage closure time. (B) Asymmetric closure, measured as the difference in the displacement of C2z and C1z, $u_2 - u_1$ , versus percentage closure time.

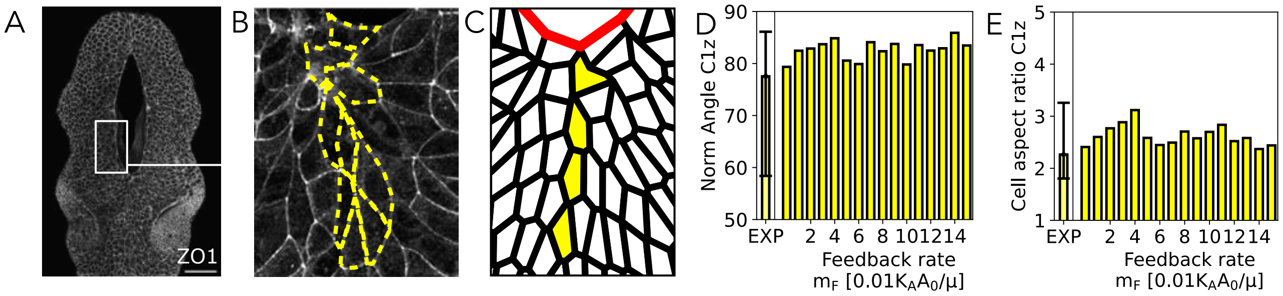


Fig. S8: **Morpho-analysis of caudal midline cells.** (A-B) HNP gap in a 14 somite fixed embryo (A), indicating caudal midline cells in yellow (B). 100 µm. (C) Zoomed in snapshot of the midline cells situated just below C1z at 60% closure. Midline region is defined by a rectangle of height equals $10\sqrt{A_0}$ and width equals $1\sqrt{A_0}$, right below the zippering point. (D,E) Comparison of midline cell orientation (D) and elongation (E) as a function of the feedback rate $m_{\mathrm{F}}$, between experiments (62 individual cells taken from 8 fixed embryos at 15-17 somite stage), and simulations. Bars denote median value, while the error bars denote the 25/75 percentile range. Simulation data account for information during the time window when row 1 consists of 20 to 50 cells, mirroring experimental conditions. The norm angle is characterized by a maximum value of 90°, indicating alignment with the rostro-caudal axis.

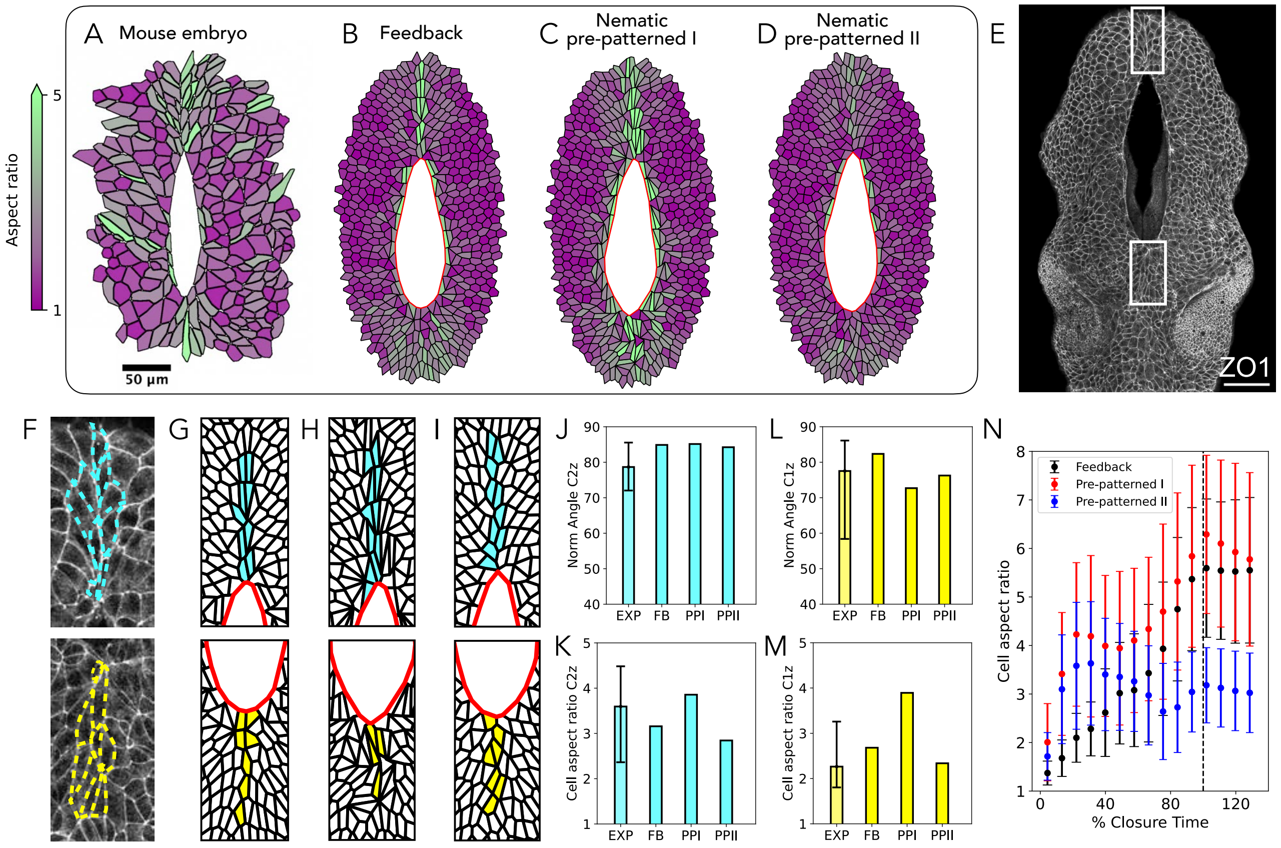


Fig. S9: **Cell patterning comparison between mouse embryos and different models with mechanical feedback or pre-patterned stress.** (A-D) Surface ectoderm cell shape patterning in (A) a 15 somite mouse embryo, and different computational simulations at 70% of HNP closure. (B) Feedback model described in the main text. (C) Nematic pre-patterned I: at $t = 0$, row-1 cells instantaneously generate a nematic organization whose principal axis follows the gap contour, with $|\mathbf{Q}_\alpha| = 1$. Cells keep fixed their nematic order parameter $\mathbf{Q}$ along the entire simulation. (D) Nematic pre-patterned II: at $t = 0$, row-1 cells instantaneously generate a nematic organization whose principal axis follows the gap contour, with $|\mathbf{Q}_\alpha| = 1$. Cells change their nematic order parameter $\mathbf{Q}$ via alignment and decay. (E) HNP gap in a 14 somite fixed embryo. Scale bar, 100 µm. (F) Zoom-in white rectangles shown in (E), indicating midline cells situated just above C2z (green) and below C1z (magenta). (G-I) Zoom-in snapshots of the midline cells at 70% closure, for the (G) feedback, (H) Nematic pre-patterned I, and (I) Nematic pre-patterned II model, respectively. Midline region is defined by a rectangle of height equals $10\sqrt{A_0}$ and width equals $1\sqrt{A_0}$, right above and below the gap. (J,K) Quantification of the median cell orientation and aspect ratio for midline cells above C2z. (L,M) Quantification of the medial cell orientation and aspect ratio for midline cells below C1z. Bars denote median value, while the error bars denote the 25/75 percentile range. (N) Comparison of the mean aspect ratio over time for cells that become midline cells (above and below the gap) between 30% and 90% of closure. Error bars represent $\pm 1$ standard deviation.

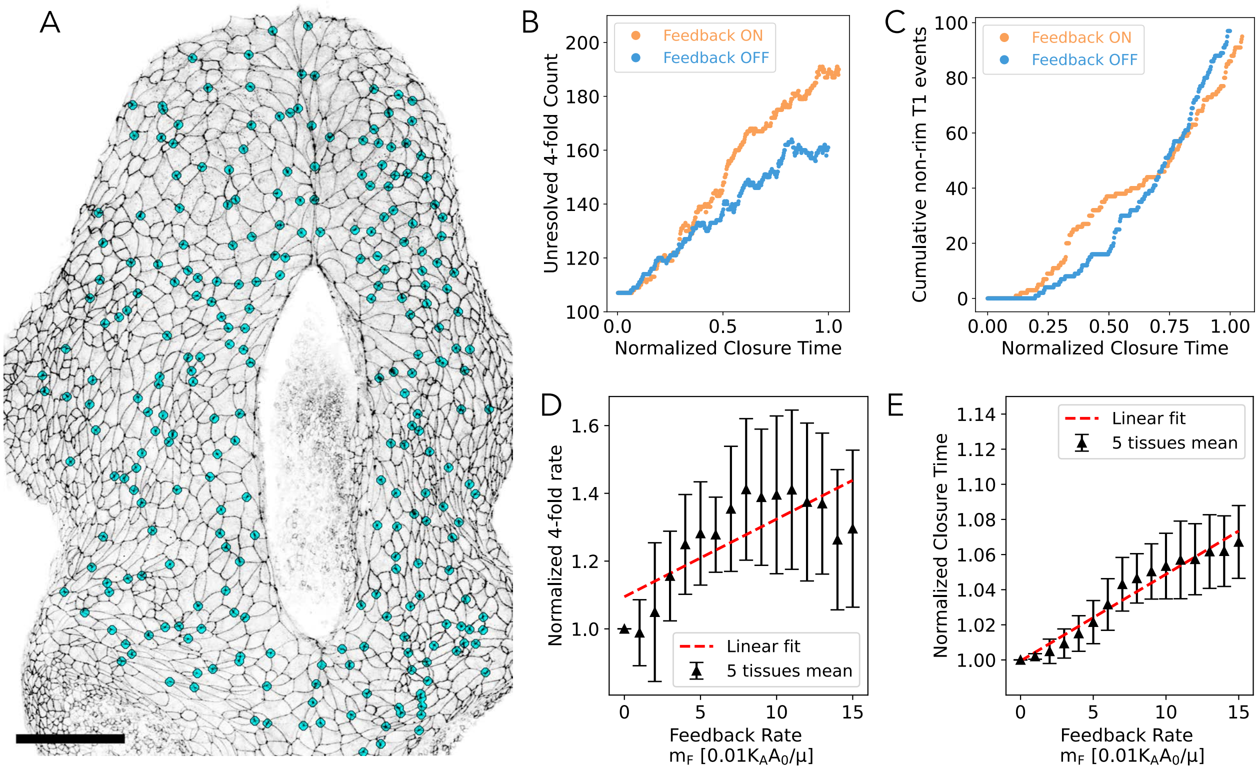


Fig. S10: **Tissue solidification during HNP closure.** (A) HNP gap in a fixed embryo, with cyan dots indicating higher-order vertices (4-fold and higher) present in the surface ectoderm. Scale bar, 100 µm. (B) Absolute count of 4-fold vertices present at any given normalized closure time in simulations ($m_\mathrm{F} = 0.08$) with and without feedback ($m_\mathrm{F} = 0$). (C) Comparison of the bulk cumulative T1 events over time, with ($m_\mathrm{F} = 0.08$) and without ($m_\mathrm{F} = 0$) feedback. We exclude the T1 events at the inner and outer boundary. (D-E) Analysis of the feedback dependent tissue solidification considering 5 simulations (geometric noise) per value of $m_\mathrm{F}$. (D) Normalized 4-fold vertex rate, defined as the slope of the count of 4-fold vertices presents in the system over time, between 20% and 80% of closure. (E) Normalized closure time. Each data point indicate the mean taken between 5 different simulations (varying initial conditions). Error bars represent ±1 standard deviation.

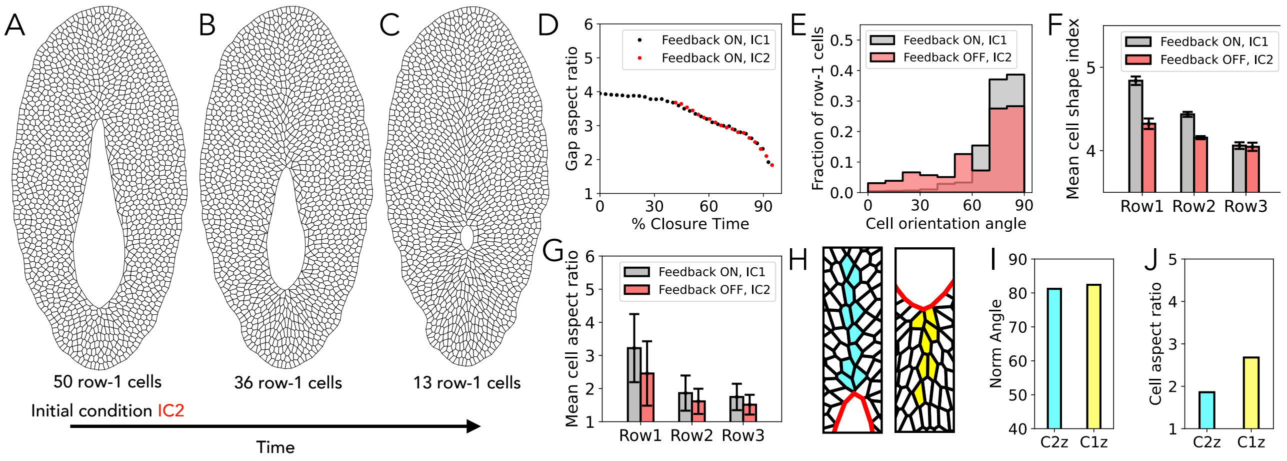


Fig. S11: **Comparison of different initial conditions and presence of feedback mechanism during simulated HNP closure.** (A–C) Tissue snapshots of an HNP closure process in the absence of feedback, initialized from the initial condition IC2, with 50 row-1 cells elongated along the gap (A). This initial condition is obtained by stopping the default HNP closure simulation with feedback defined by $\Sigma_0/K_P = 1$ and $m_\mathrm{F} = 0.08$ at 40% closure, and allowing the system to relax its mechanical energy while fixing the vertices that form the gap boundary. (B–C) Two distinct time points with 36 and 13 row-1 cells, respectively. (D–G) Gap and cell-level analysis comparing the default simulation with feedback (defined by $\Sigma_0/K_P = 1$ and $m_\mathrm{F} = 0.08$, using the default initial condition, IC1) versus the closure simulation with IC2 in the absence of feedback. (D) HNP gap aspect ratio ($H/W$) versus percentage closure time. (E) Row-1 cellular orientation. (F) Mean cell shape index per row. (G) Mean cell aspect ratio per row. The cell shape analyses in (E–G) exclude cells whose centers lie above C2z and below C1z, and consider the time window when row-1 consists of 13–36 cells, comparable to the analyses of embryos. Error bars represent $\pm 1$ standard deviation. (H) Zoom-in snapshot of the midline cells at the stage shown in (B). (I-J) Quantification of the median cell orientation (I) and aspect ratio (J) for midline cells. Analysis shown in (I-J) consider the time window when row-1 consists of 20-50 cells, comparable to the analyses of embryos.

### 5 Video Legends

**Movie S1: 60min live-imaging of a 14 somite stage mouse embryo showing HNP closure.** C2z is at the top and C1z at the bottom of each frame. Scale bar, 100 µm.

**Movie S2: Simulated HNP closure via cell crawling and actomyosin purse-string, in the absence of mechanical feedback.** C2z is at the top and C1z at the bottom of the movie. The red contour highlights the high-tension purse-string along the HNP rim. Cells around the lateral borders of the HNP show minimal elongation and their orientation lie preferentially perpendicular to the gap.

**Movie S3: Simulated HNP closure via cell crawling and actomyosin purse-string, in the presence of mechanical feedback ($m_\mathbf{F} = 0.08$ and $\Sigma_0/K_P = 1$).** C2z is at the top and C1z at the bottom of the movie. The red contour highlights the high-tension purse-string along the HNP rim. The cell surface color represents the magnitude of the nematic order parameter $\mathbf{Q}$, while the white bars indicate the nematic director. Cells around the lateral borders of the HNP exhibit robust rostro-caudal (C2z-C1z axis) elongation and orientation.

**Movie S4: 130min high-resolution live-imaging of mouse embryo showing higher-order vertices during HNP closure.** C2z is at the top and C1z at the bottom of the movie. Cyan dots highlight the positions of higher-order vertices across the surface ectoderm.

**Movie S5: Higher-order vertices predicted by our mechanosensitive feedback model.** Zoomed-in partial video of Movie S3. C2z is at the top and C1z at the bottom of the movie. The red contour highlights the high-tension purse-string along the HNP rim. Cyan dots indicate the positions of 4-fold vertices. Cyan cells are the same ones colored in Fig. 5C.
